# DNA Traces on the Shroud of Turin: Metagenomics of the 1978 Official Sample Collection

**DOI:** 10.64898/2026.03.19.712852

**Authors:** Gianni Barcaccia, Nicola Rambaldi Migliore, Giovanni Gabelli, Vincenzo Agostini, Fabio Palumbo, Elisabetta Moroni, Valeria Nicolini, Liangliang Gao, Grazia Mattutino, Andrew Porter, Pawel Palmowski, Noemi Procopio, Ugo A. Perego, Massimo Iorizzo, Timothy F. Sharbel, Pierluigi Baima Bollone, Antonio Torroni, Andrea Squartini, Alessandro Achilli

## Abstract

This research provides original insights into the diversity of DNA extracted from samples collected in 1978 from the Turin Shroud, revealing its biological complexity through rigorous DNA and metagenomic analyses. Our findings highlight its preservation conditions and environmental interactions, offering valuable perspectives into the identified genetic variants, which originated from multiple biological sources. Several human mitochondrial DNA (mtDNA) lineages were identified, including K1a1b1a, which matches the 1978 official collector’s mitogenome, H2a2 (*i*.*e*. the lineage of the mtDNA reference sequence rRCS), H1b, which is common in Western Eurasia, and H33, which is prevalent in the Near East and frequent among the Druze. Moreover, the reconstructed microbiome of the Shroud reveals a rich tapestry of multiple microbes commonly found on the human epidermis, as well as archaeal communities adapted to high salinity, and fungi including molds. This is indicative of the Shroud’s preservation conditions over the centuries. Additionally, the presence of abundant Mediterranean endemic red coral, various cultivated plants (*e*.*g*. carrot, wheat, corn, bananas, and peanuts) and domesticated animals (*e*.*g*. cattle, pigs, chickens, dogs, and cats) provide a fascinating glimpse into the diverse biological sources of the contaminants that have accumulated on the Turin Shroud over time. Finally, radiocarbon dating of two distinct threads collected from the reliquary provides evidence of their use to repair the Shroud in the years 1534 and 1694 of the Common Era (CE).

**Significance statement:** An in-depth metagenomic analysis was conducted on several linen strands collected from different areas of the body image of the *Man of the Shroud* during the official sampling in 1978. Our analyses revealed several human mtDNA lineages, including one common in Western Eurasia and another prevalent in the Near East. Additionally, the diversity of animal and plant species identified details the significant environmental contamination of the Shroud that likely occurred in recent centuries, particularly following the voyages of Marco Polo and Christopher Columbus. Radiocarbon dating of two distinct textile residuals from the Shroud’s reliquary indicated a time range between 1451 and 1800 CE, overlapping with the period of its repair interventions.

## Introduction

The Shroud of Turin is a linen burial cloth, approximately 4.4 meters long and 1.1 meters wide, which shows the double bodily image, front and back, of a man who has apparently suffered physical trauma through crucifixion with signs that are interpreted and believed to be compatible with those described in the New Testament (1, 2). It has been venerated for centuries, especially by members of the Catholic Church, and popular belief has this artifact as the cloth used to wrap the lifeless body of the biblical Jesus of Nazareth, who is acknowledged as the founder of the Christian movement that started in the Judean region in the mid-first century Common Era (CE).

The true nature of the Shroud has been highly debated among historians, theologians, and scientists. Several hypotheses regarding its origin have been proposed, but as of today, no one has yet been able to establish to any degree of certainty when and where this textile originated from. The only historical fact that has been recorded is that the Shroud has been in Europe since the mid-14th century, when it first appeared in Lirey, France, between 1353 and 1357. Starting in 1453, it was kept in Chambéry by the Dukes of Savoy until 1578, followed by several moves between France and the northern part of Italy during the 16th century, with the final resting place in Turin. Kept in a reliquary from 1694 to 1993, the Shroud was held permanently in the chapel adjacent to the Turin Cathedral, except for a brief move to Genoa in 1706 and to Montevergine, in the province of Avellino, during the Second World War. After a fire in the chapel in 1997, the Shroud was moved to the left chapel in the transept of the Turin Cathedral (for details, see (3–5)).

Considering available information and material, it has not yet been possible to either confirm or refute that the Shroud existed before its appearance in France in the mid-14th century. However, a commonly held belief is that its journey through the years began in Jerusalem, in a tomb in the Holy Land, approximately 2000 years ago, in the year 31 or 33 CE ((4) and references therein). Tradition narrates that after it was concealed for years, the Shroud would have been kept in what is modern-day Turkey, first in Mesopotamia then in Edessa, followed by a transfer to Şanliurfa, possibly around 200 CE. Subsequent moves would have seen the Shroud in Anatolia, Constantinople (modern-day Istanbul), in 944 CE (5). A burial cloth, which some historians believe to be the Shroud, was owned by the Byzantine emperors but disappeared during the sacking of Constantinople in 1204 CE (6). After this event, presumably the Shroud would have been taken by crusaders and transferred to Athens, Greece, where it would have remained until 1225. However, there are no official documents that support the presence of the Shroud in any of the above-mentioned locations.

Iconographic representations on Byzantine coins, the so-called gold *solidi* used during the Eastern Roman Empire, portray the face of the Man of the Shroud during the time of Justinian II, the Byzantine emperor who lived between 669 and 711 CE. Numismatic experts believe that the engraver must have seen the Shroud before creating the image shown on this coin because of some distinctive facial traits in common with the sacred relic. Among the various historical proofs, it is worth mentioning the votive medallion of Lirey, found in France, which shows the frontal and dorsal image of the Man of the Shroud, probably dated between 1356 and 1370. In 1988, the age of the linen cloth of the Shroud of Turin was assessed by accelerator mass spectrometry, and results of radiocarbon measurements from separate independent laboratories yielded a calendar age range of 1260-1390 CE, thus suggesting a recent Medieval origin (7). However, the raw data of the radiocarbon measurements were not initially released and only in 2017 were they made accessible through the British Museum. Statistical analysis of the original raw data strongly suggests an absence of homogeneity between the datasets made by the three laboratories, which may have undermined their reliability (8), thus requiring additional testing to ascertain its age.

The possible existence of the Shroud prior to the first documented information places the long journey of this artifact into a Middle or Near East geographical context, with a potential historical age preceding the Sacking of Constantinople in 1204. This would have then been followed by a Western European relocation and subsequent reappearance at Lirey in 1353 or 1356. Our current study attempts to differentiate between the two referenced periods separated by the so called “missing years”, considering the two geographical areas characterized by distinctive genomic data associated with human lineages, plant species, or/and animal breeds. Furthermore, we divide the chronological history of this artifact into “pre-1204”, which could involve any geographical area of the ancient Near East, and a plausible “post-1353” location in western Europe. These two hypothetical temporal and spatial differentiations may be reflected in the variation of DNA data obtained from the Shroud.

Genomic DNA was previously isolated and analyzed from organic residues recovered in 1978 from different parts of the body image on the back of the Shroud, and from threads taken from the edges and used in 1988 for radiocarbon dating of this sacred artifact (9). Analysis of the DNA extracted from organic particles and textile strands sampled from the Shroud revealed chloroplast and mitochondrial genome sequences representing multiple plant species, as well as various human biogeographical origins. In particular, these analyses revealed the presence of at least 19 plant species common in the Mediterranean Basin, in addition to plants whose primary center of origin is Asia (China), the Middle East, and the Americas, some of which reflects their introduction to the Old World after the 12th century. In addition, the analyses revealed sequences from at least 14 people of different geographic ancestry, based on the Eurasian haplogroups identified which are common to Western Europe and North-eastern Africa, as well as from the Middle East, the Arabian Peninsula, the Caucasus region, and also rare haplotypes from the Indian sub-continent (9). A quantitative assessment of the overall amplicon sequencing data revealed that over 55.6% of the human DNA corresponds to lineages from the Near East, while Western European lineages account for less than 5.6%. The presence of 38.7% of the overall human genomic data from Indian lineages is unexpected and is potentially linked to historical interactions associated with importing linen or yarn from regions near the Indus Valley, referred to as “Hindoyin” according to rabbinic texts (10).

The present study aimed to isolate and characterize genomic DNA sequences from Turin Shroud samples using a different molecular methodology (NGS vs. PCR) and an original sample consisting of linen filaments collected in 1978 by the late Prof. Pierluigi Baima Bollone from various sections of the Turin Shroud (11, 12). After DNA extraction in clean rooms, shotgun genomic libraries were sequenced using an NGS strategy with the main goal to determine the quantity and quality of human DNA and any residual DNA from microbial, fungal, plant, and animal species. The genetic results obtained were then correlated with historical information, the corresponding geographical areas of origin, and the modern distribution of plant species and human geographic groups with the aim of acquiring new data and providing novel clues as to the origin of the Shroud of Turin.

## Results and Discussion

### Human DNA sequences identified on the Turin Shroud

Before testing a total of twelve samples collected from the Turin Shroud (TS) (Table S1 and Figure S1), we verified the possibility of isolating and sequencing human DNA fragments from a linen cloth that had been purposely contaminated by touch. We obtained a low-coverage genome (2.5×) from the human control (lab operator; Dataset S1A).

We extracted DNA, built genomic libraries and obtained genomic reads from seven TS samples (Figure 1 and Dataset S1B). The percentage of trimmed reads that mapped to the human reference genome exceeded 10% in three cases, resulting in an average depth higher than 0.5× (higher than 1× for AB1 and B12). The average length of the reads (less than 100 base pairs) is consistent with the characteristics of degraded, possibly ancient DNA. However, both the damage patterns and the contamination tests show clear signatures of modern DNA contamination (Dataset S1B), as expected given the large number of people who have come into contact with the TS. To further characterize the possible presence of ancient DNA, we extracted, from each sample, reads showing patterns of deamination at their terminal bases. We applied the maximum-likelihood probabilistic model implemented in PMDtools (13) to assign post-mortem damage (PMD) scores to each read. The distribution of PMD scores is very similar among the seven samples, with most reads (>98.3%) showing a score lower than 1 (Figure S2). Only a small fraction showed PMDs greater than 3 (0.21% - 0.53%), which could be considered the minimum threshold for reliable ancient reads. For a threshold of PMDs > 5, between 0.05% and 0.19% of the total sequences were retained, comparable to values found in modern and historical individuals (13). An example of the damage patterns at different PMD scores is reported in the supplementary information for sample AB1 (Figure S3).

**Figure 1.**
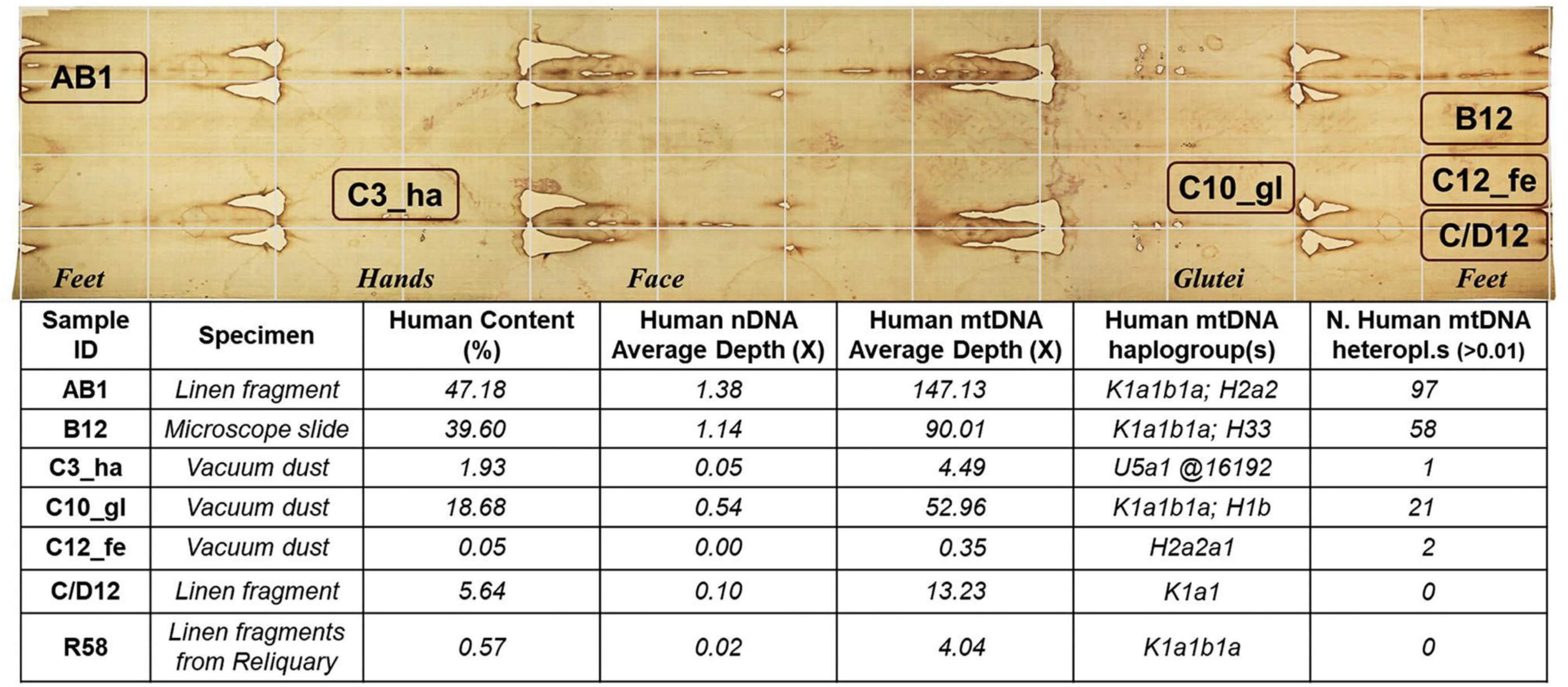
Summary of the human genomic data obtained from the TS samples. See Table S1, Figure S1, and Dataset S1 for further details.

The genetic sex of these samples was determined to be likely that of a biological male (Dataset S1B), and a predominant mitochondrial DNA (mtDNA) haplogroup was determined with high accuracy (Q>0.95; Dataset S2A) in samples AB1, B12, and C10-gl. This haplogroup, corresponding to K1a1b1a, is a typical Ashkenazi subclade of K1a1b1 with a likely Western European origin (14). The same haplogroup was identified by analyzing the mitochondrial genome of the late Prof. Baima Bollone (the collector) (Dataset S2B). Therefore, we performed kinship analyses, which revealed a close relationship (same individual or first-degree) between the three TS DNAs and the genome of the collector, who has Jewish ancestry (Table S2). As a negative control, we confirmed the complete unrelatedness between the TS genomic data (and the collector) and the modern linen specimen, artificially contaminated by the operator.

Other haplogroups were also identified in the three samples (Figure 1 and S4), namely H2a2, H33 and H1b. The first two were also reported by Barcaccia *et al*. (9). H2a2 is expected since it is the lineage of the mitochondrial reference sequence rRCS (NC_012920) and therefore can be identified in any sample without distinctive haplogroup markers. H1b is commonly found in Europe, and to a lesser degree also in Central Asia (15). H33 is a rare haplogroup found today mainly in the Near East, especially among the Druze, an Arabic-speaking ethno-religious minority currently present in the Holy Land, Jordan, Lebanon, and Syria (16, 17). Although several of the previously reported mitochondrial variants were identified among the heteroplasmies (Dataset S2A), no additional haplogroups could be identified with high accuracy. This is due to the high-quality filtering used for variant calling (*i*.*e*., 176 heteroplasmic variants are excluded from the final haplotypes) and the ability to analyze entire mitogenomes, which greatly increases the accuracy of classification, up to more than 70% (18). Our results indicate that the human DNA found on the TS specimens mostly derives from the main collector, and shows variable level of additional contamination.

### Proteomics analyses

We used liquid chromatography–tandem mass spectrometry to analyze the peptide profiles of three TS samples (namely, C/D12, B12, and R58). After excluding four proteins identified in the negatives—keratin type II cytoskeletal 1, triosephosphate isomerase, actin, and trypsin (deemed normal contaminants or deliberately added to the sample)—and other common contaminants, such as myosin and keratins 9 and 10, the only unique proteins identified in the C/D12 and R58 samples were proteinase K, with 16 unique peptides identified overall, and collagen alpha-1(I) chain, with two unique peptides identified in the R58 sample (Dataset S3). The presence of proteinase K in the samples was expected because the TS samples had been treated with proteinase K during previous attempts at DNA extraction. Since the peptides of the collagen alpha-1(I) chain showed no modification, they likely originated from skin contamination rather than from ancient peptides. This is further supported by the high presence of keratins, which are much higher in the samples than in the negative controls, indicating poor handling procedures rather than issues associated with laboratory analysis. Overall, it can be concluded that no ancient peptides were identified in the samples, only keratins and other proteins associated with skin handling.

### Radiocarbon Dating

Two linen threads collected from the reliquary were quantitatively sufficient for 14C radiocarbon dating (R58-1 and R58-2, respectively), which was outsourced to the Curt-Engelhorn-Zentrum Archäometrie (CEZA) in Mannheim, Germany (Figure S5). The obtained ages (Dataset S1C) are consistent with two historical events. The time range of sample R58-1 (1451-1622 CE) overlaps with the fire that broke out in 1532 CE in the Chapel of Saint-Chapelle in Chambéry, where the Turin Shroud was being kept inside a silver reliquary. The intense heat caused by contact with molten silver burned through the folded cloth. The Shroud was saved from destruction, and in 1534 CE, Poor Clare nuns repaired the burn holes by sewing linen patches over the damaged areas (19). Further conservation mending work was carried out in 1694 CE, likely under the direction of Sebastian Valfrè, to stabilize the Shroud (20). This work overlaps with the time range of sample R58-2 (1642-1800 CE).

### Metagenomics analyses

Due to the low number of reads from the Turin Shroud samples that mapped to the human reference genome, we conducted a comprehensive metagenomic analysis. We utilized the Kraken program for the initial screening. The distribution of reads across kingdoms varied significantly (Table S3, Dataset S5), with unassigned reads ranging from 17% (AB1) to 72% (R58). Reads assigned to viruses and archaea were minimal, while bacterial species consistently accounted for 10% to 31% of the reads. The percentages of eukaryotic reads differed significantly and showed an inverse relationship with unassigned reads; samples with fewer unassigned reads had a higher proportion of eukaryotic reads and *vice versa*. Regarding the eukaryotic reads, the vast majority were assigned to the genus *Homo*: 93% or more in AB1, C/D12, C10_gl, and B12; 86% in C3_ha; and 77% in R58 (Dataset S4). A notable exception was sample C12_fe, containing only 0.4% eukaryotic reads, of which 26% were assigned to *Homo*. As in the other samples, most of the remaining reads were assigned to Viridiplantae (e.g., 61% in the case of C12_fe).

The data derived from the Kraken classification enabled identification of Bacteria, Archaea, and Fungi, but were insufficient for Metazoa and Viridiplantae analyses due to limited sequence databases. Therefore, we adopted an alternative approach based on aligning the sequences against the NCBI nt database. Because running BLAST on all individual reads was not feasible due to their high number and short length, we performed a metagenomic assembly of the reads and subsequently aligned the resulting contigs to the database using BLAST. Challenges due to species diversity and DNA degradation resulted in low assembly metrics and short contigs (with an average length of 210-250 nt, Table S4). The results of the BLASTn analysis were categorized as those from Kraken (Table S5), showing comparable trends with generally fewer unassigned contigs due to the larger NCBI nt database used. Samples R58 and C12_fe were exceptions with higher unassigned contig shares (83.8% and 95.1%, respectively). Both archaea and viruses were present only in trace amounts. Compared to the Kraken analysis, eukaryotic contig proportions increased in all samples, likely because of assembly biases that favored the abundant *Homo sapiens* genome. A significant number of bacterial contigs could not be classified at the species or phylum levels. Only a few archaea taxa were detected (a maximum of 28 in B12 and none in R58).

The fungi kingdom was used for comparison in the analyses. Identified orders showed strong similarity across samples, although less represented orders exhibited some variability. Kraken was deemed more reliable for species quantification, while BLAST performed better for species identification despite potential false negatives.

The Kraken results were further validated through an additional analysis with MetaPhlAn, a tool specifically designed to profile microbial communities from metagenomic shotgun sequencing (21). Unlike Kraken, which uses k-mer analysis across entire genomes, MetaPhlAn aligns unpaired reads to a comprehensive microbial gene database for precise identification. Consequently, MetaPhlAn identified fewer reads, with unclassified readings from 44% (AB1) to 99% (R58 and C12_fe). The detected taxa mostly aligned with the findings derived from Kraken analysis, identifying minimal archaea and the same bacterial classes. New species constituted only 2–4% of the reads in samples R58 and C12_fe, and were attributed to *Geobacillus jurassicus*, a thermophilic bacterium (22) (Dataset S6). The concordance of the outputs obtained with two different methodologies (and reference databases) reinforces our results.

### Bacterial and Fungal taxa identified on the Turin Shroud

Overall sequencing data for bacterial, archaeal, and fungal biota, including the relative abundance of the phyla, classes, and orders, respectively, are summarized in Figure 2, with further details provided in the supplementary information (Datasets S4 and S5). Considering current knowledge of microbial ecology, the analysis of bacterial taxa associated with the TS samples revealed a notable presence of species commonly found on the human skin, particularly in sample AB1 and to a lesser extent in sample C3_ha. These include *Cutibacterium acnes* (formerly *Propionibacterium acnes*), *Staphylococcus epidermidis*, *Cutibacterium granulosum*, *Corynebacterium tuberculostearicum*, *Staphylococcus capitis*, and *Staphylococcus hominis* (23). Their presence indicates potential human skin contact with the analyzed samples.

**Figure 2.**
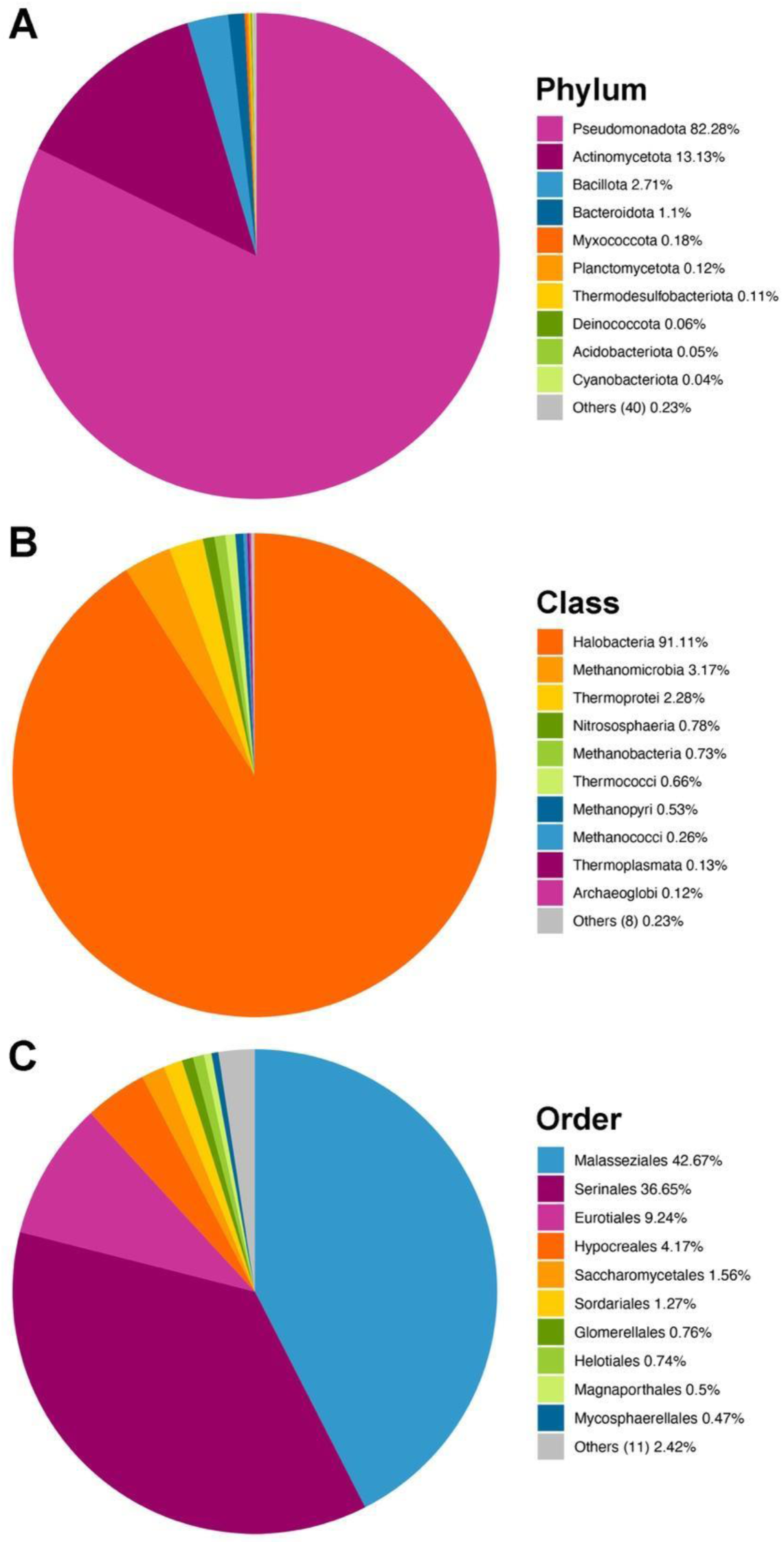
Cumulative taxonomic assignment of the TS reads. The results show the relative abundance of the bacterial phyla (A), archaeal classes (B), and fungal orders (C).

Within the archaeal domain, the TS samples exhibited strong consistency in community composition, with most taxa recurring among the top entries. These predominantly belonged to the salt-tolerant class Halobacteria, which is adapted to high salinity environments. These findings suggest a pronounced overrepresentation of osmotically resilient species, which are typically associated with saline and drought-stressed conditions.

The fungal community displayed a significant abundance of *Debaryomyces hansenii*, a yeast species that thrives in dry and osmotically stressful environments (24), similar to the environments where the Turin Shroud was kept in the past. Saprophytic molds, such as *Aspergillus,* were also identified, along with fungi associated with human skin and scalp, particularly *Malassezia restricta*.

Taxonomic identification of a diverse array of bacterial, archaeal, and fungal taxa provided insight into the complex microbiome associated with the TS, shedding light on possible ecological and historical contexts. Bacterial profiles featured prominently skin-associated taxa, which indicate direct human contact, both in ancient and modern times. The high concentration of these bacteria in sample AB1, which is from a peripheral area likely exposed to environmental contamination, suggests frequent manipulation. Sample C/D12, taken from the hand region of the Shroud’s image, further supports evidence of human contact. The detection of *Paraburkholderia* species, known for their ecological adaptability and occasional pathogenicity (25), may reflect broader environmental exposures over time.

Archaeal analysis revealed a consistent dominance of Halobacteria, which suggests an adaptation to saline environments. This could be related to ancient flax retting practices in brackish or salty water (26), which were more common than dew retting in antiquity (27). The dominance of *Debaryomyces hansenii* further supports salt tolerance and possible marine connections because the yeast is also found in seawater (24). The presence of *Aspergillus* points to possible degradation related to fluctuating storage conditions. The widespread presence of *Malassezia restricta*, a primary colonizer of the human skin (28), across all samples is a clear demonstration of ongoing human interaction with the Shroud.

Overall, the microbiological diversity of the Shroud samples offers valuable insight into its history. The recurrent presence of skin-associated bacteria and fungi implies direct contact with human beings, possibly during devotional religious ceremonies and cloth conservation treatments, in addition to the official sampling operations. These findings underscore the Shroud’s dual significance as a religious relic and a historical artifact, reflecting both devotional and environmental influences throughout its existence.

### Plant and Animal taxa identified on the Turin Shroud

A metagenomic assembly of the reads into contigs (see Material and Methods) and a nucleotide BLAST search enabled to extend the analysis to other kingdoms (Figure 3, and Dataset S7 and Dataset S8). In addition to *Homo sapiens*, our analysis of animal taxa revealed notable findings, particularly regarding the red coral (*Corallium rubrum*; 33.3%), a species endemic to the Mediterranean Sea (Figure 3B-C, and Dataset S7 and Dataset S8).

**Figure 3.**
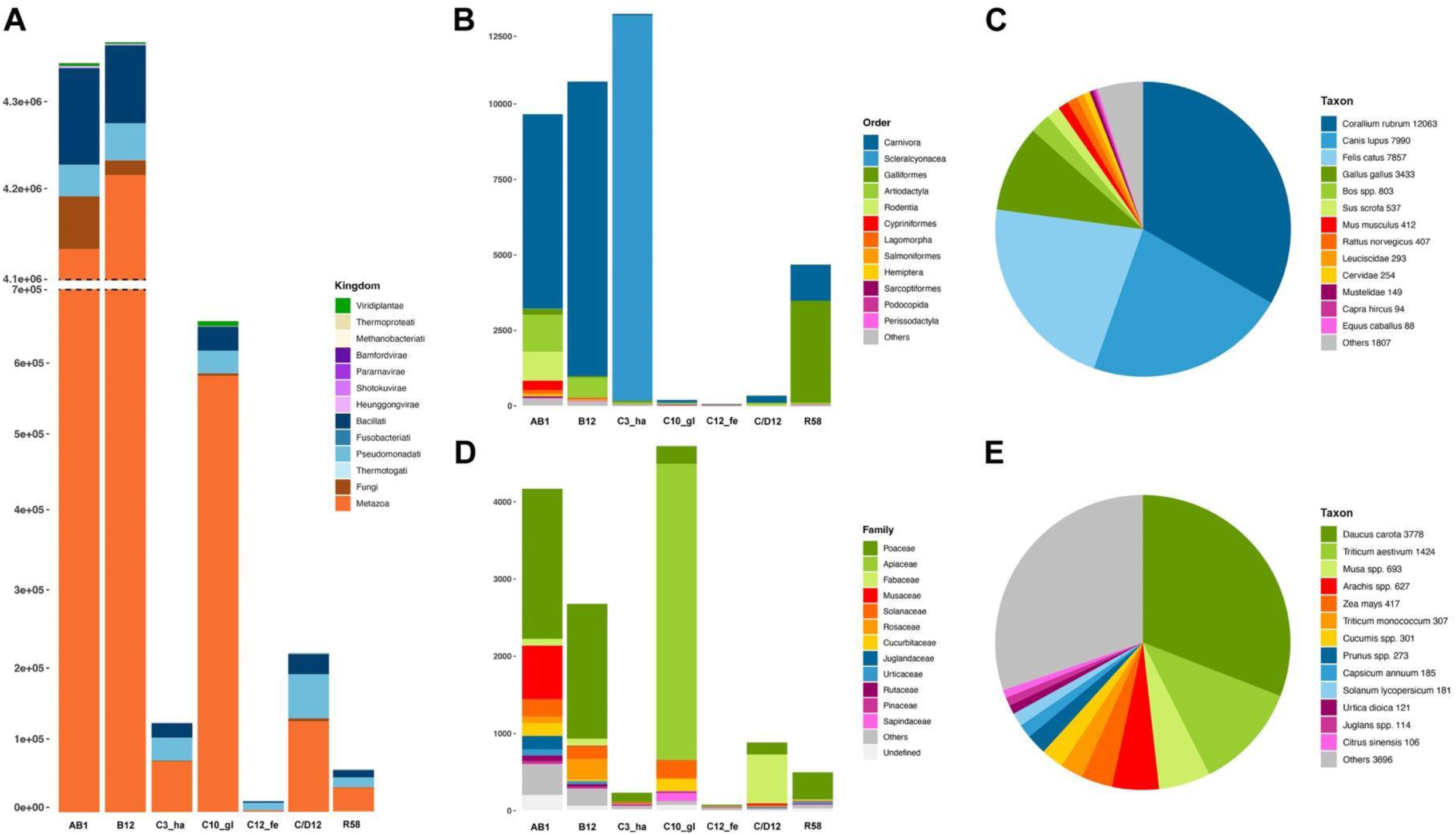
Graphical summary of the results of the taxonomic assignment of contigs. Number of contigs for each TS sample and their distribution among kingdoms (A), orders of Metazoa (B), assigned taxa of Metazoa (C), families of Viridiplantae (D), assigned taxa of Viridiplantae (E).

We extracted the reads showing patterns of deamination at their terminal bases from the animal and plant samples with more than 100 contigs, to evaluate signals of damage pattern, as in the human analysis (Figures S6 and S7).

We focused on the taxa whose identification was based on hits with at least 100 assigned contigs to avoid false positives, and found domestic cats and dogs to be prevalent (21.7% and 22.1%, respectively), although DNA traces from several farm species, including chickens (*Gallus gallus domesticus*; 9,5%) and ruminants from the Bovidae family (cattle, goats, and sheep; 2,6% overall) were detected. Other livestock and wild animals such as pigs, horses, deer, and rabbits were also found. Weak traces of brown rats and mice suggest possible environmental contamination. Among the less abundant taxa identified (around 1% of assigned contigs), notable are some fish species including the gray mullet (*Chelon labrosus*), Atlantic cod (*Gadus morhua*), and ray-finned fishes (*Leuciscinae* spp.). Marine ostracods, particularly the small seafloor-dwelling crustacean *Notodromas monachal*, which is common in zooplankton communities, were also documented. Insects such as flies (Diptera), aphids (Hemiptera), and arachnids like dust and skin mites and ticks, including *Demodex folliculorum* and *Haplochthonius simplex*, were identified as well.

Plant taxa showed greater diversity (Figure 3D-E, and Dataset S7 and Dataset S8), with prominent species including carrot (*Daucus carota*; 30.9%) and bread wheat (*Triticum aestivum*; 11.6%), represented by more than 100 assigned contigs. Other cereals found encompassed durum wheat (*Triticum durum*), einkorn wheat (*Triticum monococcum*), maize (*Zea mays*) and rye (*Secale cereale*). Horticultural crops included peppers, tomatoes, and potatoes (Solanaceae), as well as melons or cucumbers (Cucurbitaceae). The Fabaceae family was also represented by a strong presence of peanuts (*Arachis* spp.), particularly in sample C/D12. Among forage species and their wild relatives, weak traces of perennial ryegrass, bluegrass, fescue, oats, and clovers were observed. The fruit trees identified with greater confidence included banana, almond, walnut, and sweet orange. Fainter DNA signals were detected for fig, pistachio, apple, pear, hazelnut, maple, and grapevine (*Vitis* spp.). The only non-cultivated plant with more than 100 assigned contigs was stinging nettle, but significant amounts of spruce trees and wild cardoons were also found (see Dataset S7 and Dataset S8).

The taxonomic identification of the plant and animal species on the Shroud reveals a complex biodiversity, albeit constrained by limitations in available reference databases. Reliance on a relatively small dataset (126 plant species and limited animal genomes) and the degraded condition of the DNA available made the taxonomic classification complicated. Nevertheless, several Viridiplantae taxa were confidently identified, particularly agricultural species such as carrots and cereals. Grouping of taxa at the genus level (*e.g*., *Triticum*, *Aegilops*, *Hordeum*, and *Secale* spp.) reflects a parsimonious approach due to the overlapping of genetic signals. However, the strong signal for *Daucus carota* indicates that higher specificity could be obtained in some cases.

Among cultivated species, we also identified melon or cucumber, and various Solanaceae (tomato, pepper and potato). These reflect typical agricultural practices in the Mediterranean and other European territories. Fruit trees such as banana (*Musa acuminata*) and orange (*Citrus sinensis*) further highlight the diversity of cultivated plants possibly introduced through trade and indicate interconnected regional economies.

Non-cultivated species such as cardoon (*Cynara cardunculus*), maple (*Acer* spp.), spruce (*Picea* spp.), pine (*Pinus* spp.), nettle (*Urtica dioica*), and walnut (*Juglans* spp.) reflect local ecological diversity. Notably, cardoon and nettle are indigenous to Europe and the Mediterranean area. Wild cardoon remains widespread in southern Europe and North Africa, and it was domesticated in the western Mediterranean (29, 30). Stinging nettle, native to Europe and widely distributed across Asia and North Africa, is valued both as a vegetable and in traditional Indian medicine (31). The uncertain origin of melon is also noted, with hypotheses pointing to ancient Persia or Africa (32).

Although none of the identified plant taxa can be uniquely associated with the Levant and the Middle East, several genera originated in the Fertile Crescent and are now widespread across the Mediterranean Basin. For example, almond trees, pistachios, and figs are native to and have been domesticated in the Near or Middle Eastern areas (33–35), following the introduction by Romans to Southern Europe and Northern Africa during the first centuries CE.

The presence of exotic species introduced to Europe from Central and Eastern Asia, and North and South America, was also noted. Maples, for instance, are thought to have originated in Asia, with genetic studies suggesting that the genus *Acer* likely emerged in East Asia, particularly in China. However, fossil evidence indicates that some species may have evolved independently in North America (36). Bananas, whose secondary center of diversification is Africa, were introduced to Europe by Portuguese sailors in the early sixteenth century, but they gained popularity only by the eighteenth century. This shift was driven by improved agricultural practices and trade networks, including plantation systems in Central America (*e*.*g*., the Caribbean), which helped establish bananas as a staple fruit in Europe (37).

Several plant taxa identified here, such as *Picea abies*, *Juglans regia*, *Pinus* spp. *Cucumis* spp., *Cichorium* spp., *Cynara* spp., and *Vitis* spp., were also found in previous palynological and/or molecular studies (9, 38–40). These findings support a view of the Shroud as being associated with a blend of local vegetation and historical agricultural practices.

Animal taxa presented a more complex identification challenge due to the genetic similarities among related species. The presence of *Corallium rubrum* underscores the region’s historical reliance on marine resources. This endemic Mediterranean red coral has been known since the time of the Roman Empire and held considerable cultural, economic, and symbolic significance in antiquity. It was primarily fashioned into jewelry, amulets, talismans, and decorative objects. Several ancient texts from the Middle Ages attest to its use in medical contexts, as it was believed to possess healing properties and was utilized in remedies for various ailments and disorders. Regarded as a valuable commodity, red coral was transported for centuries along Mediterranean trade routes, facilitating economic and cultural exchanges among populations and regions. The trade of red coral saw further development during the medieval period, particularly in Mediterranean ports (41, 42).

Livestock species as well as cats and dogs, reflect long-standing agricultural and domestication practices in local farms and rural settings. The presence of mites and ostracods, including the abundant *Notodromas monacha*, which are very common in terrestrial and aquatic habitats illustrates the ecological diversity and may reflect environmental conditions during the storage or handling of the Shroud.

### Genetic affinities among the Turin Shroud samples

To highlight potential similarities among the TS samples, we performed two different analyses based on their metagenomic profiles. The artificially contaminated linen fragment was used as an outgroup.

The Principal Component Analysis (PCA) revealed the outlier behavior of AB1, which represents the TS later edge, while samples derived from the areas of face, hands, gluteal region, and feet of the body image of the Man of the Shroud did not show distinct patterns related to sampling type (linen fragment vs. vacuum dust) (Figure 4A). On the other hand, C12_fe and R58, collected from the Shroud and the Reliquary, respectively, were the samples with the highest number of private taxa in the hierarchical clustering plot (Figure 4B). It is important to note that the outgroup used as control was confirmed as outlier by both analyses and its genetic differentiation from each of the TS samples was comparable. In addition, although the first two components accounted for about 72% of the overall genetic variation, most TS samples were closely clustered by PCA, except for AB1. This low discrimination is partially due to the method, which could obscure differences between samples (see Materials and Methods). However, a closer examination of the overall genetic data (Dataset S7) reveals the presence of both different and similar organisms among the pairwise comparisons of the TS samples. These include several domesticated species, such as cereals, vegetables, pets, and farm animals. We believe that this commonality can be explained by two hypotheses rather than solely by the sample history. The first hypothesis is technical: these species are well-described and included in various databases, thus any misalignment during identification is likely to occur with higher frequency. The other hypothesis calls for secondary contamination, which refers to DNA traces resulting not from the direct exposure of the TS samples to those particular species, but rather from human contact with those organisms. Basically, we could hypothesize the presence of a sort of secondary metagenomic fingerprint. This would lead to a collection of species that appear with greater probability than others. However, this hypothesis could not be explored further within our project because it would require a comparison of additional samples (similar to the TS) from different origins.

**Figure 4.**
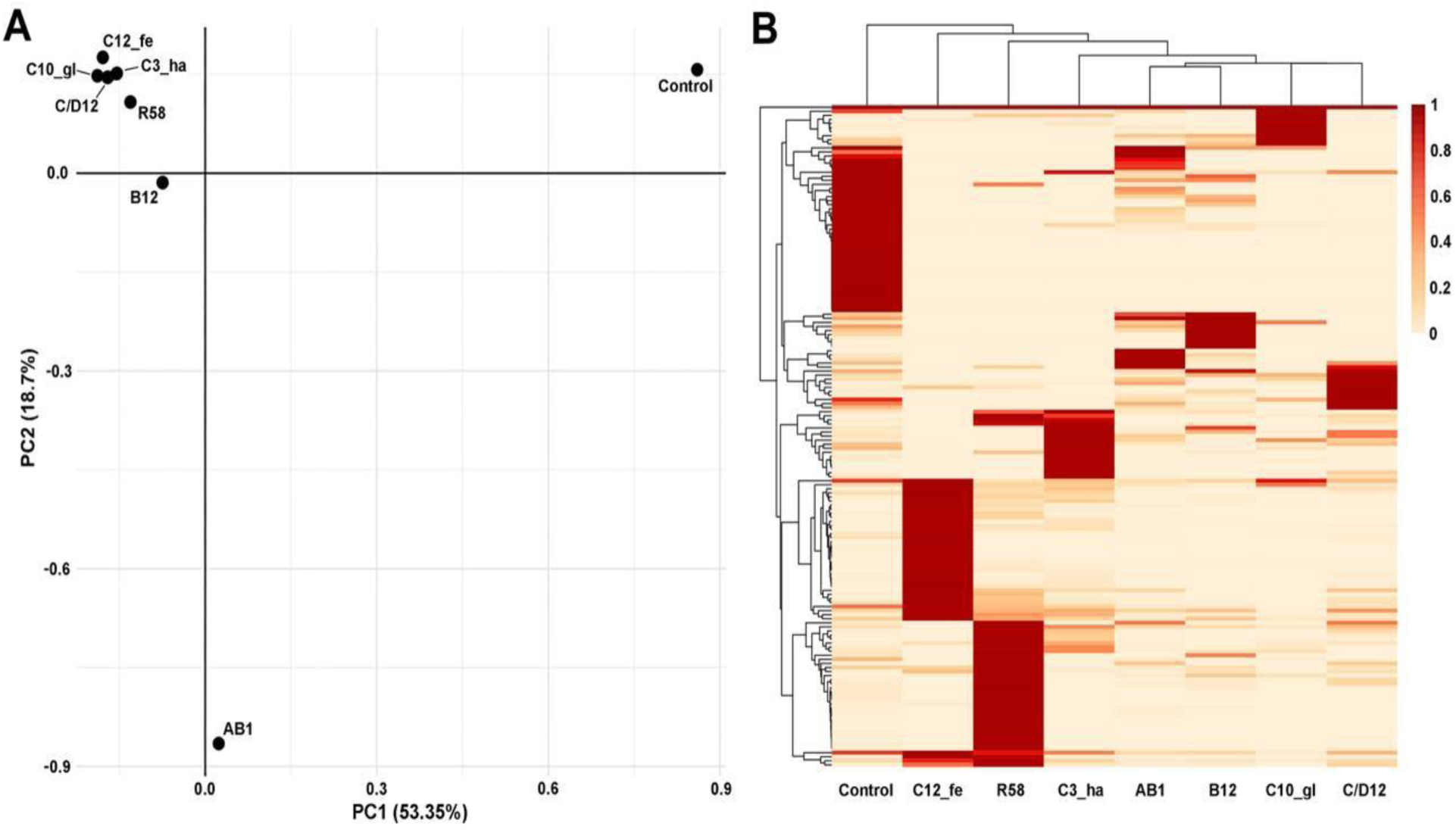
Clustering analyses. (A) Principal Component Analysis and (B) Hierarchical Clustering based on BLASTn metagenomic data of the seven TS samples and an artificially contaminated linen fragment (control).

The presence of recurrent species, while potentially introducing background noise, must be emphasized: numerous other species were identified exclusively in specific samples of the Shroud or with exceptional clarity, such as *Daucus carota* and *Corallum rubrum*.

### Reappraisal of the overall biodiversity found on the Turin Shroud

The study aimed to isolate and characterize genomic DNA sequences from the available original official samples of the Turin Shroud collected in 1978 by Prof. Pierluigi Baima Bollone from various sections of this linen cloth. Its final goal was to identify the biological sources of environmental contaminants—including pollen, plant fibers, fungal structures, animal hairs, blood residues, and human biological fluids—found in linen filaments, a vacuum dust sample, and a small cloth fragment. Our new findings, alongside previous ones (9), confirm the presence of plant and animal DNA, as well as traces of DNA from various human subjects on the Shroud of Turin. The identification of diverse DNA sources provided insights into potential connections between the temporal and geographic pathways related to hypotheses on the Shroud’s origin. Many identified plant species are native to and widespread in Central Europe and the Mediterranean Basin, extending from the Iberian Peninsula to the Middle East.

The most prominent crop identified in the TS samples is carrot (30.9%). Both historical records and molecular analyses suggest that carrots, characterized by purple and yellow roots, were domesticated in Central and Minor Asia, and then introduced to Western Europe (43–45). It is worth mentioning that out of approximately 3,800 contigs, about half were confidently mapped to unique sites of the carrot genome, whereas the remaining half mapped to multiple locations being likely associated with repetitive sequences. The majority of the contigs originated from the C10 sample, comprising 1,787 contigs (94.4%), whereas only 2 and 1 contigs are from AB1 and B12 samples, respectively. SNPs common to a DNA sequence dataset of carrot accessions (46) belonging to five distinct populations were employed to calculate their similarity. Based on this analysis, carrot DNA contaminations found on the Shroud were shown to be genetically more similar to improved cultivars and early cultivars (Figure 5), which are descendants of orange carrots developed in Western Europe between the 15th and 16th centuries (47). These genetic signatures suggest a relatively recent contamination – not before the late Middle Ages – due to carrot residuals most likely associated to plant materials of Western European origin.

**Figure 5.**
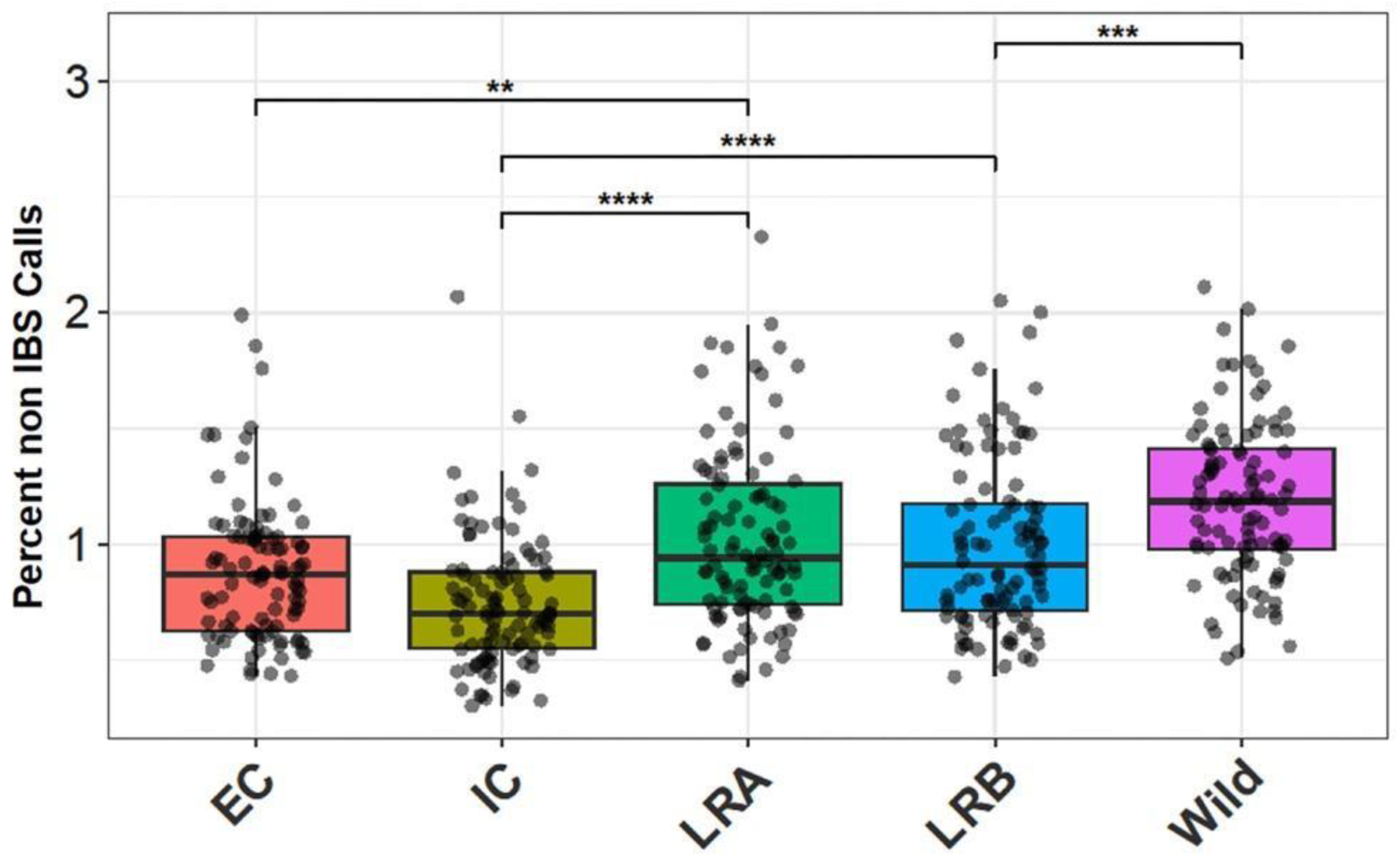
Percent of non-Identical By State (IBS) calls between contigs assembled from C10_gl sample and carrot population groups. Carrot samples represent the following populations as determined by Coe et al. (46): Early cultivars (EC), Improve cultivars (IC), Landrace A, Landrace B and Wild. Each data point represents a random subset of 20 contigs sampled from 1,787 contigs. Significant differences were tested using T-Test. Asterisks represent the following significant levels: *=0.05, **=0.01, ***=0.001, ****=0.0001.

Another abundant crop among those identified in the TS samples is wheat and its close relatives. All cultivated polyploid forms originated and were domesticated in the Fertile Crescent and were subsequently introduced in the Mediterranean Basin from the Middle East. By the 1st century BC, with the emerging Roman Empire, wheat had become well-established in the region and recognized as an essential agricultural commodity. Additionally, the presence of exotic species introduced to Europe from Central and South-Eastern Asia, and North and South America, was noted. For example, maples, oranges and bananas were identified. Maples are thought to have originated in Eastern Asia, despite some evidence suggesting a North American origin (36). Moreover, oranges, which are originally from the region encompassing Southern China and North-Eastern India, and bananas, whose earliest evidence of domestication is Papua New Guinea, were introduced to the Mediterranean Basin by the Muslim conquerors and Portuguese sailors, respectively, between the 9th and 10th centuries and 15th and 16th centuries (37, 47). The identification of animal taxa posed a more intricate challenge owing to genetic similarities among closely related species (*i.e*. members of the Bovidae family). The presence of both wild and farm animals, including several livestock species and pets, indicates the enduring nature of agricultural and domestic practices prevalent in rural settings. The diversity of animals and plants at multiple taxonomic levels suggests that much of the environmental Shroud’s contamination occurred in recent centuries, particularly after the voyages of Marco Polo and Christopher Columbus. This hypothesis is consistent with and further corroborates our previous findings (3, 9).

In this study, we also identified human mitochondrial variants, which can be classified into one main haplogroup (K1a1b1a, which is the same of the collector), and those with diagnostic motifs of different lineages, such as H1b, an ancient late glacial marker common in Western Eurasia, and H33, which is typical of the Near East and particularly frequent among the Druze (17) as previously documented by Barcaccia *et al*. (9). Notably, the Druze population shares a common genetic ancestry with Jews and Cypriots, and has historically intermingled with other Levantine populations, including Palestinians and Syrians (16, 17). Although additional single nucleotide polymorphisms (SNPs) were identified, it was not possible to reconstruct complete mitogenome haplotypes that could be confidently classified into any of the South Asian minor lineages (e.g., M39, M56, R7, and R8) identified in our previous study by analyzing mtDNA hypervariable segment I (HVS-I) from a different sample of the Shroud of Turin (9). In that study, over 55.6% of the human mtDNA HVS-I reads were classified into Near Eastern lineages, while Western European lineages accounted for less than 5.6%. The presence of approximately 38.7% of Indian ethnic lineages could have resulted from historical interactions or the Romans importing linen from regions near the Indus Valley, associated with the term “Hindoyin” found in rabbinic texts (10). Notably, the term “Shroud”, derived from the Greek “Sindôn” meaning fine linen, may be related to Sindh, a region renowned for its high-quality textiles. Historical evidence supports trade links between India and the Mediterranean, underscoring the significance of these textiles and inviting further exploration of ancient cultural interactions and trade practices. Indeed, the biblical scholar Levergne stated that the term “Sindôn” refers to a fabric of Indian origin, valued for its qualities and used for various and multiple purposes (48). In brief, a reappraisal of those outcomes from the analysis of the DNA traces found on the Shroud of Turin suggests the potentially extensive exposure of the cloth in the Mediterranean region and the possibility that the yarn was produced in India.

Overall, our prior and present findings provide valuable insights into the geographic origins of individuals who interacted with the Shroud throughout its historical journey across various regions, populations, and eras.

## Conclusions

In this study, we sequenced and characterized genomic DNA traces extracted from the cloth edges, previously used for radiocarbon dating, and from other sections bearing the image of the Man of the Shroud. Molecular traces of contamination from individuals who handled the Shroud with limited protective precautions, including the 1978 sampling team in Turin, were expected. Our analysis revealed that the predominant mitochondrial lineage is characteristic of Ashkenazi Jews and matches the mtDNA haplotype (and genomic data) of Prof. Baima Bollone, the collector, as also confirmed by kinship analyses on the entire genome. Diagnostic variants of other haplogroups, such as H33 found in Middle Eastern populations (*i.e*. Druze), were also detected, alongside the Western Eurasian haplogroups H2a2 and H1b. The occurrence of an unusually high number of human heteroplasmies and the coexistence of different mtDNA variants confirm that the Shroud came into contact with multiple individuals, thereby challenging the possibility of identifying the original DNA of the Shroud.

Intense human handling is confirmed by skin bacteria such as *Cutibacterium* and *Staphylococcus*. Halophilic archaea indicate salt preservation or saline storage conditions, but the presence of *Debaryomyces hansenii* and other fungi found in cheeses and hyper-saline waters raises further questions. Concerning the Shroud’s journey, the presence of red coral, livestock (*e.g*. chickens, cows, pigs, goats, sheep, rabbits, horses), and domestic cats and dogs suggest Mediterranean origins or transit through Mediterranean regions.

Plant DNA analyses revealed the predominance of cultivated species, which has historical implications. The concomitant and very abundant presence of both *Daucus carota* (domesticated carrot) and various *Triticum* species (cultivated wheat) suggests agricultural practices with both native and introduced crops. Carrot is the most prominent crop plant species identified in the Shroud. Wild populations of carrots with white roots have existed in Europe since more than two thousand years, as they were described by Pliny the Elder in the encyclopedic “Naturalis Historia” and used by Romans for medicinal purposes and food preparations (see Apicius, “De re culinaria” a collection of Roman cookery recipes). However, we demonstrated that the carrot DNA found on the Shroud is genetically more similar to early cultivars and improved cultivars, which were proven to descend from orange carrot varieties developed in Europe between the 15th and 16th centuries (46). Furthermore, oranges and bananas, which are originally from Southern Asia and Eastern Oceania (China and Papua New Guinea), were introduced to the Mediterranean Basin by Muslim populations and Portuguese sailors in the high-late Middle Ages (37, 47). Post-Columbus species, such as maize (domesticated in Mexico), Solanaceae crops (*e.g*. tomatoes, peppers, and potatoes), and peanuts (Fabaceae), which are crop plants native to Latin America, pose an additional significant conundrum about the antiquity of the Shroud, even if a later contamination cannot be ruled out. DNA traces from various species and regions, including the Middle East, Mediterranean, Europe, America and Asia, indicate that the Shroud was exposed to different environments and peoples. The prevalence of Mediterranean crops and the absence of typical Middle Eastern flora raise questions about the agricultural landscape when the Shroud was created or used as a burial cloth. Comparative analyses with other ancient textiles and artifacts could further illuminate cultural and historical interactions with plants and animals.

In conclusion, genetic and microbial evidence discloses a complex history of the Turin Shroud, reflecting interactions with a diverse array of individuals and exposure to various agricultural and environmental contexts. While some human mtDNA haplogroups may correspond to those who handled and sampled the historical artifact, the overall DNA results – derived from rigorous methodological handling in clean rooms and metagenomic analyses supported by robust bioinformatics – suggest a diverse mosaic of genetic traces. The age of the Turin Shroud cannot be determined through metagenomics because this methodology is unable to provide any robust evidence supporting either a Medieval origin or a history dating back two millennia. Nevertheless, our findings constitute a novel and significant contribution to the field, thoroughly elucidating the biological traces left by centuries of social, cultural, and ecological engagement. Furthermore, the radiocarbon dates of two distinct fabric threads collected from the reliquary confirm their use in mending and patching efforts aimed at stabilizing the fire-damaged linen cloth during two well-documented events in 1534 and 1694 CE. Collectively, our findings illuminate important aspects of the Shroud’s preservation history.

### Addendum

In memory of Dr. Pierluigi Baima Bollone, coauthor of this publication, whose sudden departure has left our research groups at a loss for words.

As Pierluigi Baima Bollone passed away in Turin, Italy, on November 5, 2025, we wish to express our profound gratitude for his irreplaceable collaboration, invaluable contribution and unwavering dedication to the advancement of knowledge regarding the Turin Shroud.

Bibliographical notes: Pierluigi Baima Bollone was a prominent medical doctor specialized in hematopathology and a full professor of forensic medicine at the University of Turin. He founded and led the Department of Diagnosis and Prevention at Gradenigo Hospital. A prolific author of scientific publications, dissemination books and forensic medicine manuals, his works are widely adopted in academia. Renowned on both national and international stages, he produced influential writings in criminology and forensic medicine. He also served as Honorary President of the International Centre for Sindonology in Turin, the official institution devoted to the study of the Shroud, contributing extensively to its literature.

## Materials and Methods

### Sample collection

For the present study, twelve samples were retrieved from the 1978 Turin Shroud collection (Table 1 and Figure S1): seven glass tubes with linen cloth filaments, four jars with vacuum dust (one completely empty), and one microscope slide with a tiny biological fragment. This collection is entirely new compared to the one analyzed in Barcaccia *et al*., 2015 (9).

### Sequence data production

DNA was extracted from all available samples using a silica-based technique that allows for recovery of ancient/historical DNA even when molecules are very few and highly fragmented, as in (49). After DNA extraction, DNA concentration and fragmentation were assessed by fluorometer and capillary electrophoresis, respectively. For samples with a DNA concentration higher than zero, shotgun genomic libraries were constructed following a custom double indexing protocol optimized for ancient DNA using partial uracil DNA glycosylase treatments (half-UDG), which partially retain the deaminated cytosines resulting from DNA damage (50).

Considering that 10M reads are required for an accurate estimation of the endogenous content, the half-UDG genome libraries were multiplexed and sequenced on the Illumina platforms (NextSeq 550 and 2000) using a paired-end strategy. A present-day filament of linen was artificially contaminated by one of the handlers (operator) to validate the feasibility of our approach.

The modern DNA of the collector (Prof. Baima Bollone) was also extracted using the Maxwell® RSC Stabilized Saliva DNA Kit after collecting a biological sample with the IsoHelix GeneFiX™ Saliva DNA Collection Kit. For this modern sample, a genomic library was prepared with the Illumina DNA Prep Kit using the standard protocol and sequenced on the Illumina NextSeq 2000.

### Sequence data analyses

The raw shotgun sequence data have been demultiplexed using bcl-convert v. 00.000.000.4.2.4 (51). Raw fastq and adapter trimming have been performed using AdapterRemoval v. 2.3.2 (52), discarding reads shorter than 30 bases after trimming and collapsing paired-end reads overlapping by at least 11 bases (parameters: *–trimns –maxns 2 –trimqualities –minquality 2 –minlength 30 – collapse –minalignmentlength 11*). Trimmed reads were mapped to the human reference genome hs37d5 using bwa v. 0.7.17-r1188 (53) with the aln/samse algorithms and parameters (*-l 1024 -n 0.01 -o 2*) optimized for ancient DNA (54). After mapping, aligned reads were filtered for a minimum mapping quality of 20 and duplicates were removed using samtools v. 1.17 (55) and DeDup v. 0.12.8 (56), respectively. The two terminal bases of each read in the final BAM files were soft-clipped using trimBam from BamUtil v. 1.0.15 (57), to mask potential aDNA damage patterns for downstream analyses. Statistics and aDNA damage patterns were evaluated with QualiMap v.2.2.2-dev (*bamqc*) (58) and mapDamage v. 2.1.0 (59). Genetic sex was evaluated following the two methods described in Skoglund *et al*. (60) and and Mittnik *et al*. (61). For the mtDNA, the trimmed reads were aligned to a modified version of the rCRS (62), obtained using CircularGenerator v. 1.0 from CircularMapper (63), with bwa aln/samse and the same parameters as indicated above. RealignSAMFile v. 1.0 (from CircularMapper) was used to realign the reads to the original rCRS. BAM files were filtered in the same way as above. Mitochondrial DNA haplogroups were called using Haplocheck v. 1.3.2 with default options (64).

For a reliable haplogroup classification, ANGSD v 0.940-dirty (65) was also used to call a consensus mitochondrial sequence with the following parameters: *-minQ 20-setMinDepth 2-minMapQ 30-doFasta 2-doCounts 1-basesperLine 60*. Consensus files were uploaded on HaploGrep v 3.2.1 (66). The mitogenome data were also analyzed with Haplocheck v. 1.3.2 (64), which was also used to construct schematic phylogenetic trees.

### Contamination estimate

Contamination estimate on the mtDNA was obtained using contamMix v. 1.0-10 (67). A consensus sequence from the aligned mtDNA reads was called with ANGSD v. 0.940-dirty (65) using the following parameters: *--minMapQ 30-minQ 20 -doCounts 1 -setMinDepth 3 -doFasta 2*. The original mt-aligned reads were mapped to this consensus and used, together with a database containing 311 worldwide mitochondrial genomes, plus the consensus generated with ANGSD, to estimate potential mt contamination with contamMix (parameters: *–nIter 50000 – nChains 3 –trimBases 2*). Nuclear contamination was estimated using hapConX (68) for those samples classified as male individuals.

PMDtools (13) was used to assign PMD score to all reads mapped to the human genome and to extract endogenous ancient DNA sequences, separating them from contaminating modern sequences using postmortem degradation patterns.

### Modern reference individual

The raw reads from the modern reference individual were processed using fastp v. 0.24.0 (69) with parameters *-g -x -c*. Trimmed reads were mapped to the hs37d5 human reference using bwa-mem2 (70). Duplicates were removed using picard-tools MarkDuplicates (71) and final statistics were calculated using mosdepth v. 0.3.6 with default parameters (72).

### Kinship analysis

Kinship analyses were performed using the samples showing a sufficient average depth of coverage on the human genome, together with the modern reference individual. Three different analyses were performed using READ v. 2 (73) and BREADR (74) and KIN (75). PileupCaller (from sequenceTools v. 1.5.3.2) (76) was used to call pseudohaploid genotype data on the 1240K SNPs set to be used as input data for READ2 and BREADR.

### Proteomics methods

Three TS samples (C/D12, B12, and R58) were extracted by adding 5% SDS / 50 mM TEAB buffer (20 µL to samples B12 and C/D12 and to two negative controls; 30 µL to R58). Extraction proceeded by alternating 30 min in a sonicator at 56 °C with 30 min on a thermomixer hotplate at 1,600 rpm, repeated over 5 h. Samples and negatives were then incubated overnight at 56 °C without shaking. The following day, five 1.4 mm ceramic beads, pre-cleaned with bleach, rinsed with methanol, and UV-irradiated overnight, were added to each tube. Mechanical homogenization was performed using a Precellys tissue lyser at 6,500 rpm for 30 s. Lysates were centrifuged at 12,700 rpm (max speed) and supernatants transferred to protein low-bind tubes. For each sample, a final volume of 23 µL was prepared (adding extraction buffer as needed) for downstream S-Trap processing. Reduction was carried out by adding 1 µL of 60 mM DTT and incubating at 55 °C for 15 min. Alkylation followed by adding iodoacetamide (IAM) to a final concentration of 500 mM stock solution, incubating in the dark for 30 min at room temperature. Excess IAM was quenched by adding 1 µL DTT. To acidify the solution, 2.6 µL of 27.5% phosphoric acid was added, followed by 165 µL of S-Trap binding buffer consisting of 100 mM TEAB in 90% methanol. Acidified samples were loaded onto S-Trap micro spin columns and centrifuged at 4,000 ×g for 30 s. Each column was washed three times with 150 µL of binding buffer. Digestion was performed by adding 0.5 µg sequencing-grade trypsin in 50 mM TEAB directly onto the columns, followed by incubation for 2 h at 47 °C.

Peptides were eluted following the standard three-step S-Trap elution protocol: 40 µL of 50 mM TEAB, 40 µL of 0.2% formic acid in water and 40 µL of 50% acetonitrile / 0.2% formic acid. Each elution step included centrifugation at 4,000 ×g for 30 s, and eluates were pooled. Peptide solutions were dried overnight in a fume hood prior to MS analysis.

Peptide samples were analyzed by liquid chromatography–tandem mass spectrometry (LC–MS/MS) using a Thermo Scientific Orbitrap Astral mass spectrometer coupled to a Thermo Scientific Neo nano-UHPLC system via a nano-electrospray ionization source. Peptides were separated on a reversed-phase C18 analytical column (75 µm internal diameter, 25 cm length) operated at a nominal flow rate of 300 nL/min. Mobile phase A consisted of water and mobile phase B of acetonitrile, both containing 0.1% formic acid. Peptides were eluted using a linear gradient from 1% to 55% solvent B over 45 min, followed by a column wash at 99% B and re-equilibration, for a total run time of 52 min.

The mass spectrometer was operated in positive ion mode. Full MS survey scans were acquired in the Orbitrap analyzer over an m/z range of 350–1500 at a resolution of 240,000. Data-dependent MS/MS acquisition was performed with precursor ions selected for fragmentation using an isolation window of 1.5 m/z. Fragment ions were acquired in the Astral analyzer over an m/z range of 150–1800 using higher-energy collisional dissociation (HCD). Dynamic exclusion was enabled to minimize repeated sequencing of the same precursor ions. All other instrument parameters were set according to manufacturer recommendations and are reported in the supplementary methods.

The raw data were analyzed using Progenesis QI for Proteomics version 4.2. Specifically, one experiment was conducted on the three samples only, and one separate experiment was conducted on the two negative controls, to identify proteins present in either the samples or in the negatives. After importing the data, all samples aligned against the selected reference with a similarity score greater than 90%. Peptide ions with charge 2-5 were kept, together with those with more than 2 isotopes. Overall, 3900 peptide ions were retained for the “samples” group, and 1359 for the “negatives” group. Peptides with a ranking 1-3 were exported in Mascot for identification. Search parameters were the following: semiTrypsin searches, with 2 missed cleavages allowed, fixed modification Carbamidomethyl (C) and variable modifications Deamidation (NQ) and Oxidation (M), peptide mass tolerance 10 ppm, fragment mass tolerance 0.02 Da, target FDR 1%. Results were further refined using Percolator. Peptide identified with a score smaller than 7 (for “samples”) and 12 (for “negatives”) were excluded from further analyses (as per Mascot peptide score distribution to indicate identity or extensive homology), as well as proteins identified by less than 2 unique peptides.

### Taxonomic classification of the DNA sequences

The sequenced reads from the seven samples that underwent metagenomic analyses (Table 1) were processed to trim adapters and filter for quality using Trim Galore v0.6.7 (77). Initial taxonomic classification of the reads was performed using Kraken v2.1.2 (78) with default parameters for paired-end library reads. The database used was a combination of pre-built Kraken databases, named archaea, bacteria, plasmids, human, fungi, viruses, protozoa, and plants. An interactive visualization of the results, known as a Krona graph, was generated from the textual Kraken reports using KrakenTools (see Supplementary information, including Dataset S5) (79).

An additional classification of the reads, applicable only to microorganisms, was performed using the MetaPhlAn program (21). All reads assigned by Kraken to a taxon ID belonging to Archaea, Bacteria, or Fungi, or not assigned to any taxa, were extracted using TaxonKit v0.18.0 (80), SeqKit v2.1.0 (81), and a custom awk command. The resulting paired-end FASTQ files were analyzed with MetaPhlAn v4.2.2 using the default settings, except for the parameter -t, which was set to “rel_ab_w_read_stats”.

Due to the absence of Metazoa records (except for *Homo sapiens*) in the Kraken database and the small number of Viridiplantae species included, Kraken2 analysis is not suitable for accurately identifying reads from these kingdoms. Therefore, a different approach was chosen. Raw reads were assembled into contigs for each sample using Megahit v1.2.9 (82) with the parameters --presets meta-large and --min-contig-len 100. Taxonomic classification of the resulting contigs was conducted using BLASTn v2.12.0 (83) against the entire NCBI nt database (84), downloaded on December 6, 2024. The alignment parameters were: -evalue 1e-5, -max_target_seqs 5, -perc_identity 90, and -outfmt 6. The parameters qcov, qcovhsp, and staxids, were added to the standard fields for subsequent filtering and processing.

A preliminary filtering of the alignment results was performed to generate summary statistics. Alignments with query coverage below 90% were excluded, and for each contig, the best target based on E-value was selected. A custom Bash and Python3 script, utilizing TaxonKit v0.18.0 (80), was used to calculate the number of contigs assigned to different taxa at each taxonomic rank.

The tabular output of the BLASTn alignments was further processed for taxonomic classification. For each contig, alignments with query coverage below 90% were excluded. Remaining alignments were compared based on their highest-scoring HSP (high-scoring pair) for each target. This comparison aimed to determine whether the alignments were of similar significance. Alignments were deemed non-equivalent if any of the following criteria were not met: a maximum difference in percent query coverage of 3, a maximum difference percent identity of 3, a maximum difference in bit score of 10%, a maximum difference in E-value of three orders of magnitude. If one alignment outperformed all others, the contig was assigned to the corresponding taxonomic ID (taxid). If two or more alignments were equally probable, the contig was assigned to the lowest taxonomic node present in the genealogy of all target taxid. Contigs assigned to taxa above the family rank were discarded as too unspecific. This processing was implemented using a custom Bash and R script, incorporating TaxonKit (80) and the R package rentrez v1.2.3 (85). All custom scripts are available upon request.

A final step was implemented to filter potential false positives. A manual interpretation of the tables containing all contigs assigned to a taxon was necessary to eliminate possible false positives. Two criteria were established: i) When multiple species within the same genus produced similar hits, or if the number of contigs assigned to the genus exceeded that of contigs assigned to any individual species within it, the contigs were assigned to an unidentified species within that genus (*e.g*., *Musa* spp.); ii) In cases where a substantial number of contigs identified a specific species, as well as the corresponding genus, subtribe, and tribe, all contigs were assigned to the species (*e.g*., *Daucus carota*). These reads likely represent broader, less specific regions that remain associated with the species, and it is prudent to accurately specify and categorize them accordingly.

This methodology was developed in response to findings related to monkeys, which likely yielded results associated with human DNA. Given our confidence regarding the presence of human DNA, it is highly improbable that various exotic monkey DNAs are present. We anticipate that in other instances, a small number of contigs will also identify species related to the primary target. This phenomenon may arise from the short contigs generated from degraded sequences, which can align, albeit infrequently, to incorrect targets when conserved regions are present.

### Clustering of the samples

The analysis conducted on the seven samples was also applied to the modern reference individual (hereafter referred to as “control”), with minor variations in the steps involving Fastp, Megahit, and Kraken2. The same parameters were utilized, except for the use of single-end mode instead of paired-end mode, which was dictated by the sample’s nature.

To elucidate the differences among the samples, it was imperative to select a taxonomic rank that would minimize potential assignment errors between species or genera, thereby ensuring that reads assigned to that taxon could be regarded as accurate and reliable. The variation in classification practices across different kingdoms necessitated the choice of distinct taxonomic ranks corresponding to each kingdom: Archaea at the class level, Bacteria at the phylum level, and Metazoa and Fungi at the order level, while Viridiplantae was classified at the family level.

A matrix was constructed to display the number of contigs assigned to each defined taxonomic unit across the seven samples and the control. Subsequently, a principal component analysis (PCA) was performed using the R function “prcomp”, with the parameter scale = TRUE to prevent the Primates order from disproportionately influencing the variability. This clustering effectively represents the variance among the samples. However, it has the potential to overemphasize the significance of taxa represented by only a single contig in any given sample. To mitigate this issue, a second clustering analysis was conducted, wherein all taxa with a total number of contigs less than or equal to three were excluded. The number of contigs for each sample was then normalized by dividing each value by the maximum observed value, effectively rescaling all values to a range of 0 to 1 (max normalization by columns). This was followed by a row-wise max normalization to ensure that every taxon held equal weight in the analysis. Finally, the Euclidean distances between samples were calculated and subjected to hierarchical clustering using the R functions from the pheatmap package (86).

### Genetic ancestry of carrot DNA contamination

To determine the genetic ancestry of the carrot DNA found on the Shroud, a comparative genomic analysis was performed against the *Daucus carota* reference genome (46). Contigs from Shroud samples R58, AB1, B12, C10_gl, and C/D12 — identified as *D. carota* — were mapped independently to the reference genome. In parallel, Illumina reads (∼10x) from 200 diverse carrot samples were aligned to the reference, and SNPs were identified using the GATK (Genome Analysis Toolkit) pipeline (87). The carrot samples represented five distinct populations as defined by Coe et al. (46): Early cultivars, Improved cultivars, Landrace A, Landrace B, and Wild. For each population, 40 accessions with an ancestry coefficient >0.7 were selected. Both early and improved cultivars represent lineages descended from carrot cultivars developed in the 15th and 16th centuries. Landrace A and Landrace B consist of descendants from Central/Eastern Asia and Western/Southern Asia, respectively, exhibiting semi-domesticated phenotypes. Wild accessions represent undomesticated carrots from Europe. Because samples R58, AB1, B12, and C/D12 yielded <3 uniquely mapped contigs, they were excluded from further analysis. In contrast, sample C10_gl provided 1,787 uniquely mapped contigs and was used for all downstream analyses.

To estimate genetic similarity, Identical by State (IBS) counts were calculated using the overlapping SNPs between the carrot accessions and the C10_gl contigs. To ensure the reliability of the analysis and account for potential biases in contig distribution, a subsampling approach was employed: twenty contigs were randomly selected from the 1,787 available for C10_gl, with this process repeated for 100 iterations. The resulting non-IBS percentages were analyzed using t-tests and linear models in R (88) to determine if the genetic distances between C10_gl and the known phylogenetic groups were statistically significant. In this framework, lower non-IBS percentages denote higher genetic similarity, indicating a closer ancestral relationship. Plots were generated in R using ggplot2 (89).

## Acknowledgments

The authors wish to express their gratitude to Emanuela Marinelli, an internationally recognized scholar of the Shroud of Turin, for her valuable insights and constructive comments on the sindonological aspects, particularly during the initial phase of research planning and the final phase of discussing the overall findings. Her support made this project possible and greatly enriched our work. We would also like to thank Alberto Cenzato, who assisted us in revising the Kraken code and managing the resources necessary to run it. We are deeply grateful for his work and dedication, which enabled us to overcome this technical challenge.

This study was funded by the Italian Ministry for Universities and Research (MUR) PRIN 2022 grant 2022Y8BSAL (A.A.); Fondazione Cariplo—Bando Giovani Ricercatori 2023, rif: 2023–1373 (N.R.M.); University of Padova (UNIPD), BIRD 2025, project A metagenomic survey of the linen cloth samples officially collected from the Turin Shroud in 1978, DAFNAE1–DOR–00719 (G.B.). UKRI FLF grant n. MR/Y019989/1 (N.P.). We acknowledge the CINECA award under the ISCRA initiative, for the availability of high-performance computing resources and support.

## Data availability

Genomic sequences (raw data in FASTQ format) have been deposited in the European Nucleotide Archive (ENA; https://www.ebi.ac.uk/ena/browser/home) under the accession number PRJEB88938. Any additional information required to reanalyze the data reported in this paper is available from the lead contacts upon request.

