## Supplemental figures and tables for "DNA Traces on the Shroud of Turin: Metagenomics of the 1978 Official Sample Collection"

**This PDF file includes:**

Limitations of the study
Figures S1 to S7
Tables S1 to S5
Legends for Datasets S1 to S8

**Other supporting materials for this manuscript include the following:**

Datasets S1 to S8

##### *Limitations of the study*

We encountered methodological challenges in terms of data production due to the very limited amount of source material and significant DNA degradation. Our initial goal was to verify the presence of human blood traces on the shroud fibers. Next, we examined the possible presence (and characteristics) of DNA reads of human origin. Finally, we determined the number of operational taxonomic units (OTUs) derived from environmental and individual contaminants associated with all the organisms that have come into contact with this shroud over the centuries. Finally, we tried to correlate the obtained results with available historical information, the geographic regions of probable origin, and the modern distribution of the species.

This study also faced significant challenges in data analysis, primarily due to the short length of the reads and the limitations of the reference DNA databases. These two factors likely contributed to the lower assembly metrics observed in the metagenome reconstruction and the difficulties encountered in identifying taxonomic entities univocally, particularly regarding animals and plants. These technical difficulties were compounded by the peculiarity of the proposed objectives: assigning minimal amounts of genetic material to any of the known species. These challenges required the adoption of stringent filters for species identification. Therefore, it is possible that some false negatives are present. Although relying on short contigs is advantageous for detecting minimal traces of genetic material, resulted in the alignment of many contigs in genomic regions largely conserved among taxa, eventually reducing the number of informative sequences.

#### Figures

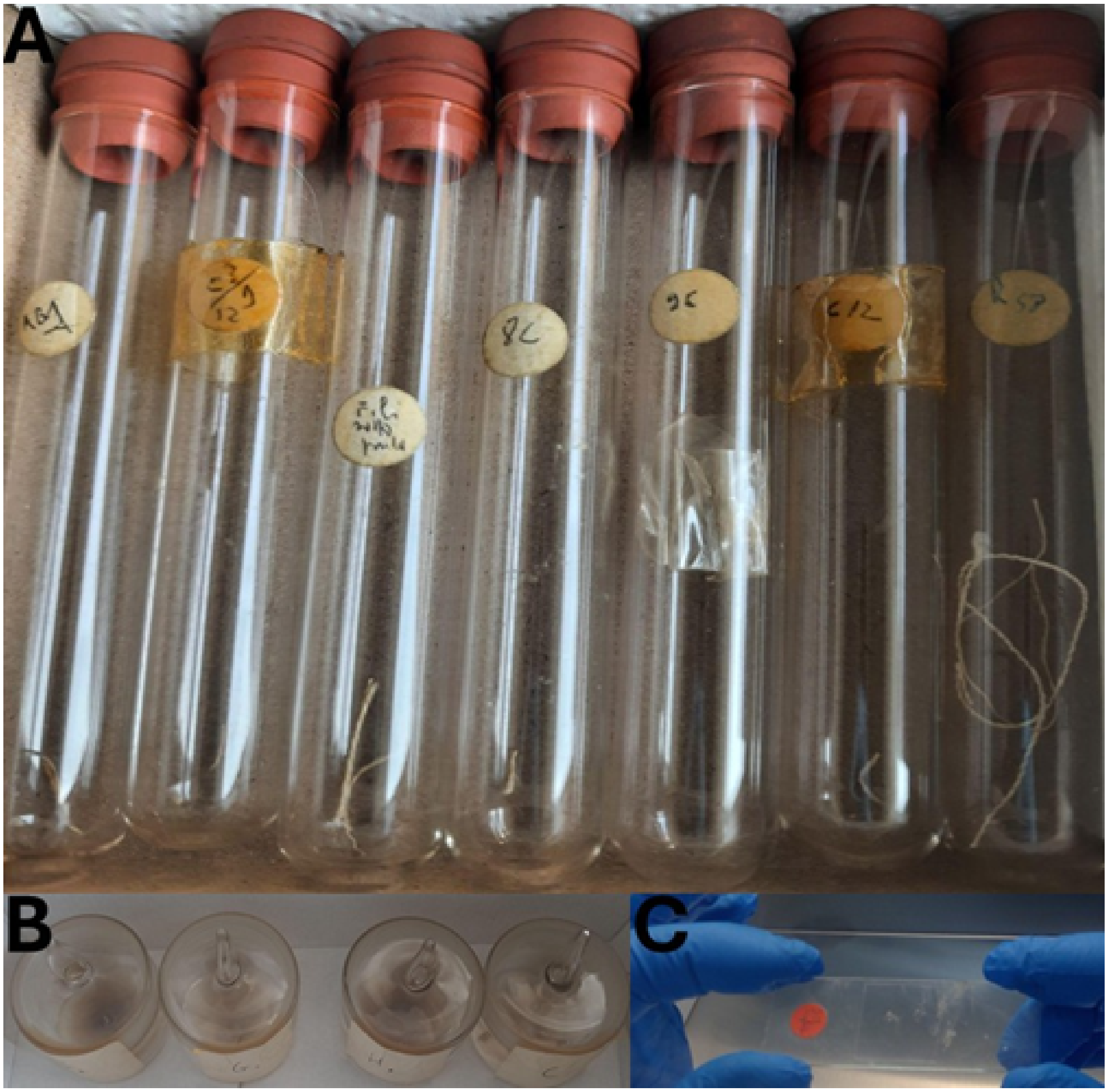


**Figure S1. Pictures of the TS samples provided by Prof. Baima Bollone (collector) and analyzed in this study.**
A) Linen cloth fragments conserved in glass tubes, see also Supplementary Table 1. The sample “*fili sotto piede*” (literally “threads under foot”) has been renamed as “A11”, while the tube labeled as “R58” refers to linen threads collected from the Reliquary (Prof. Baima Bollone report). B) Vacuum dust preserved in glass jars. C) Small biological sample on a microscope slide with a cover slip. See Table S1 for additional information.

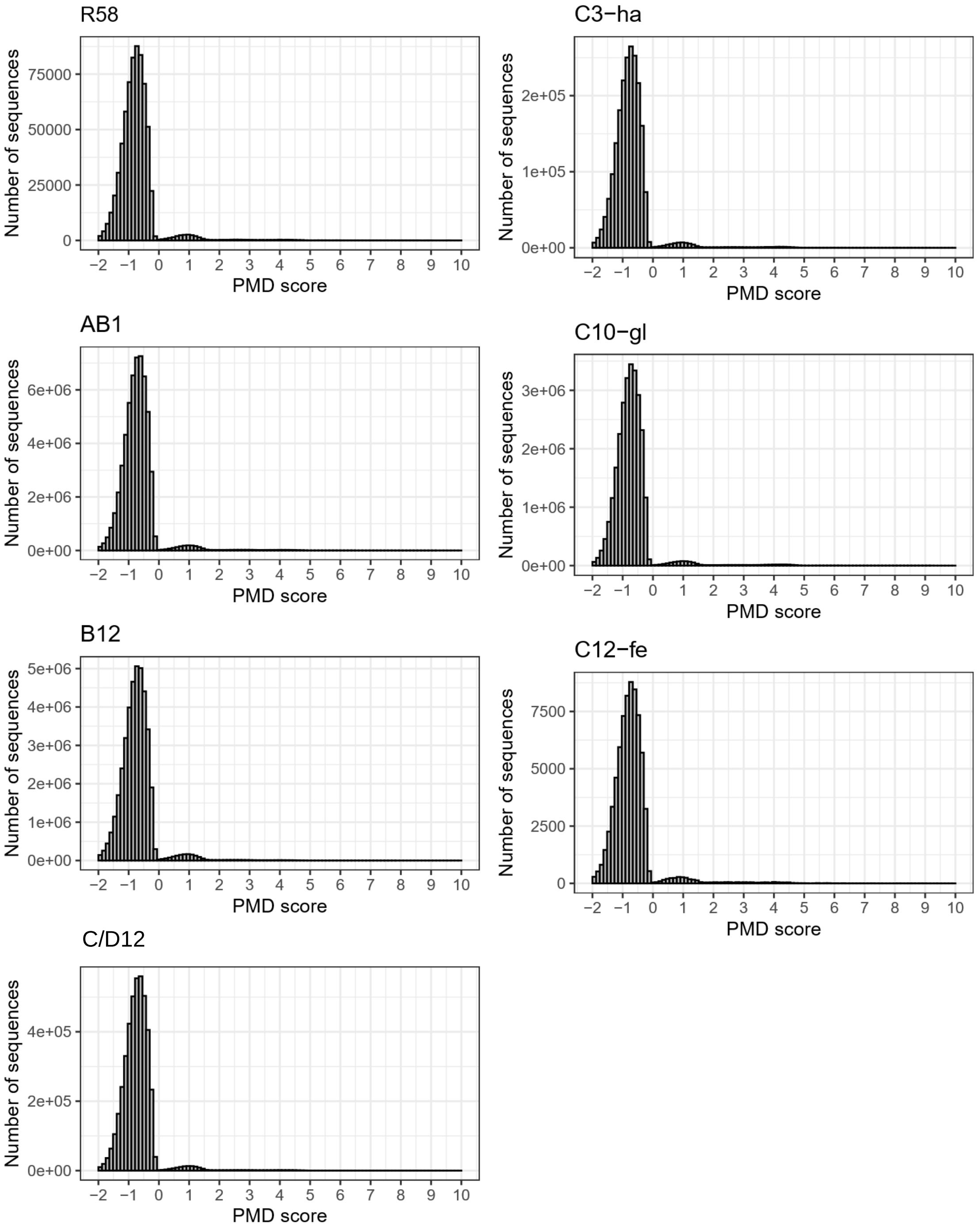


**Figure S2. PMDs distribution in the seven TS samples aligned to the human reference genome.**

The maximum-likelihood probabilistic model implemented in PMDtools [Skoglund et al. (https://doi.org/10.1073/pnas.1318934111) was applied to assign post-mortem damage (PMD) scores to each read.

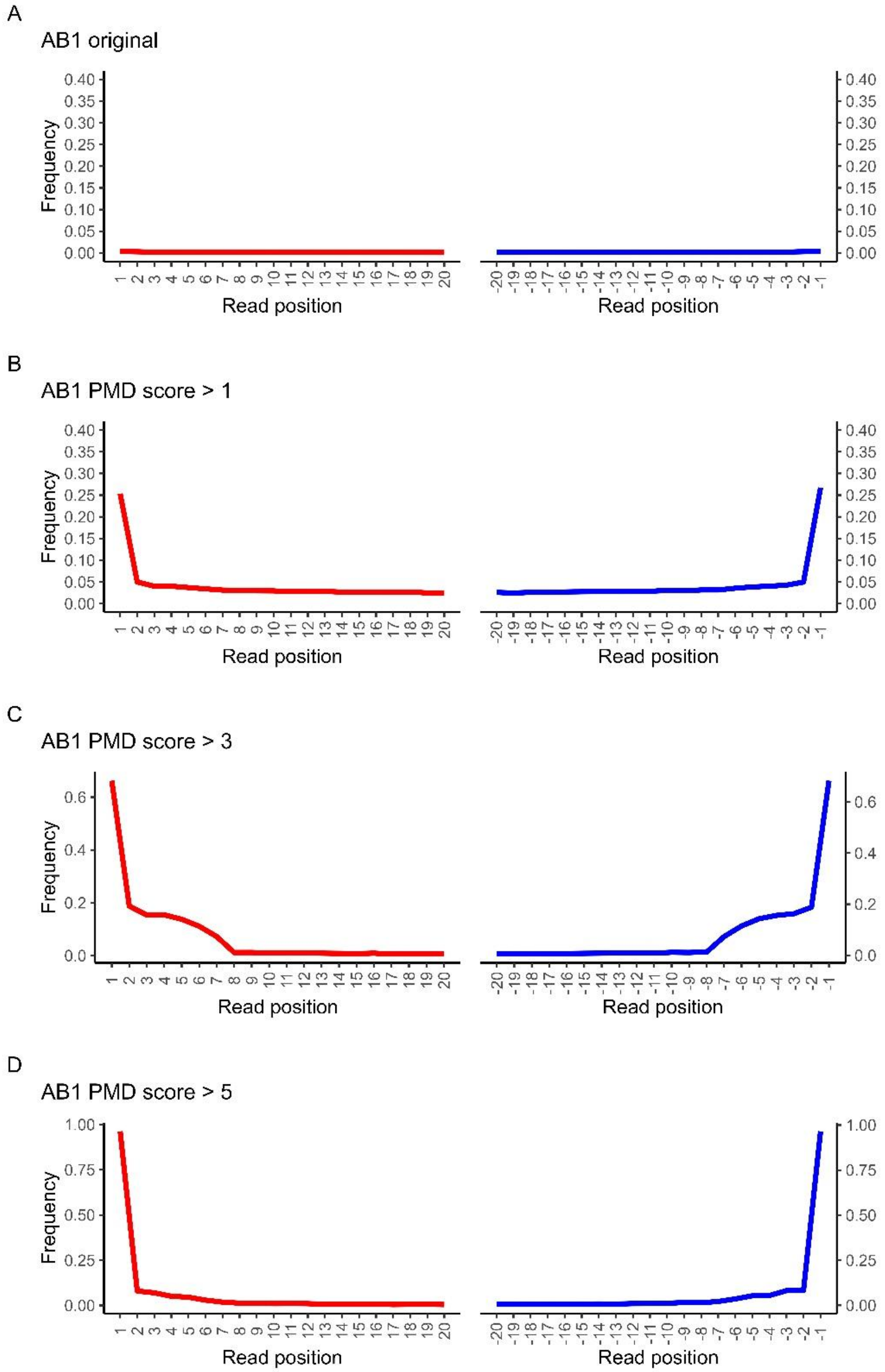


**Figure S3. Damage pattern analysis of sample AB1 mapped to the human reference genome.**

The damage pattern analysis was computed on A) the total mapped reads, B) the reads showing a PMDS greater than 1 (1.14% of the total), C) the reads showing a PMDS greater than 3 (0.25% of the total), D) the reads showing a PMDS greater than 5 (0.076% of the total).

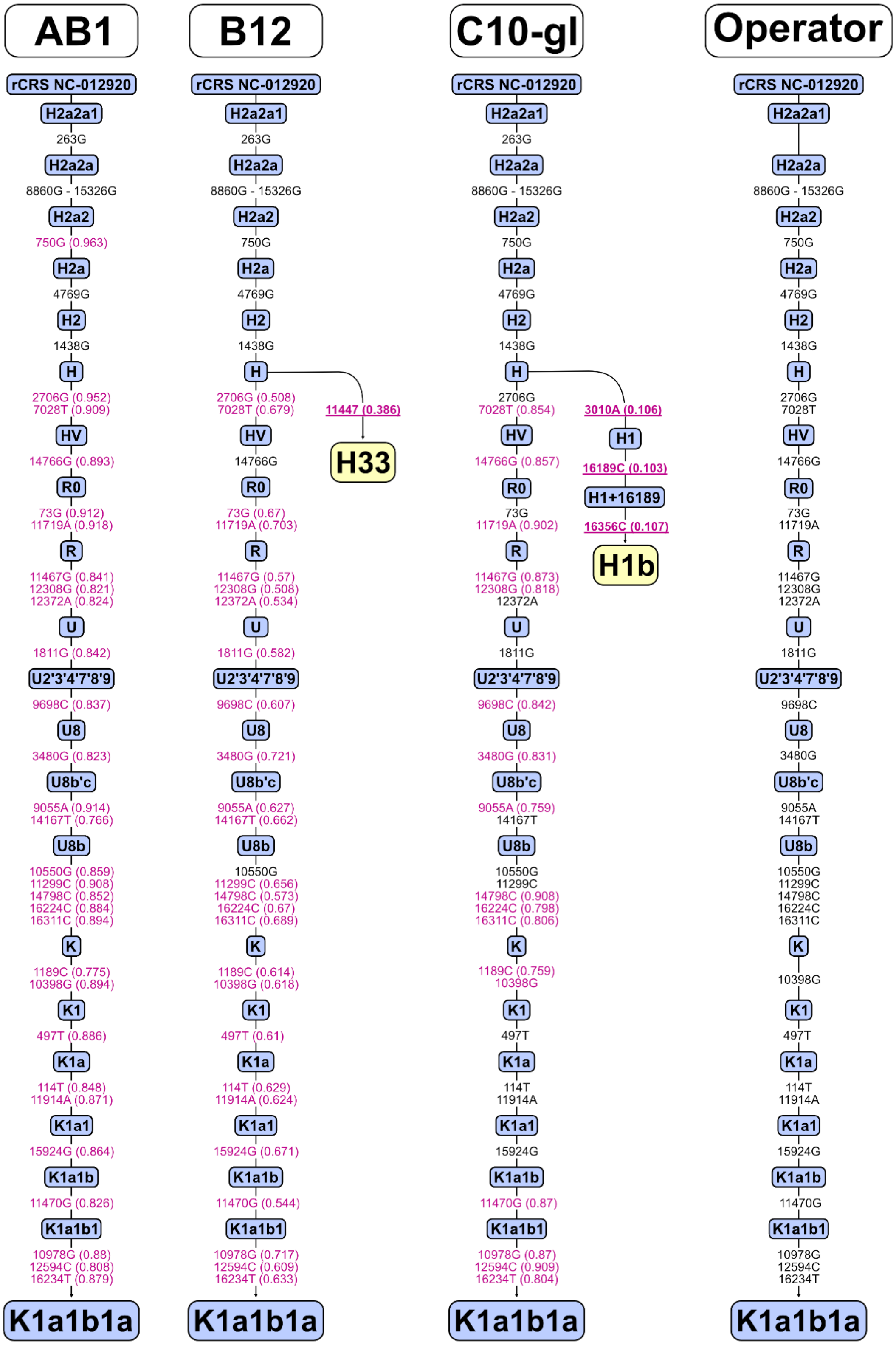


**Figure S4. Schematic phylogenetic tree and haplogroup classification of complete mitogenome data from three TS samples and the collector.**
SNP markers are homoplasmic (in black) or heteroplasmic (in violet). Numbers in parentheses indicate the frequency of each heteroplasmic variant.

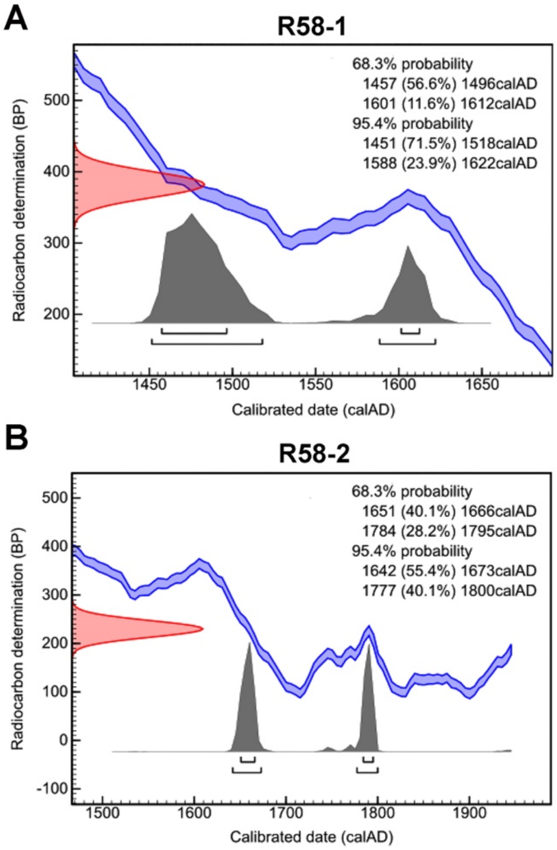


**Figure S5. Radiocarbon dating of two TS linen threads (R58-1 and R58-2) collected from the reliquary.**
The estimated ages correspond to the range 1451-1622 CE for sample R58-1 (A) and 1642-1800 CE for sample R58-2 (A). This 14C dating analysis was outsourced to the Curt-Engelhorn-Zentrum Archäometrie (CEZA) in Mannheim, Germany (see also Dataset S1C).

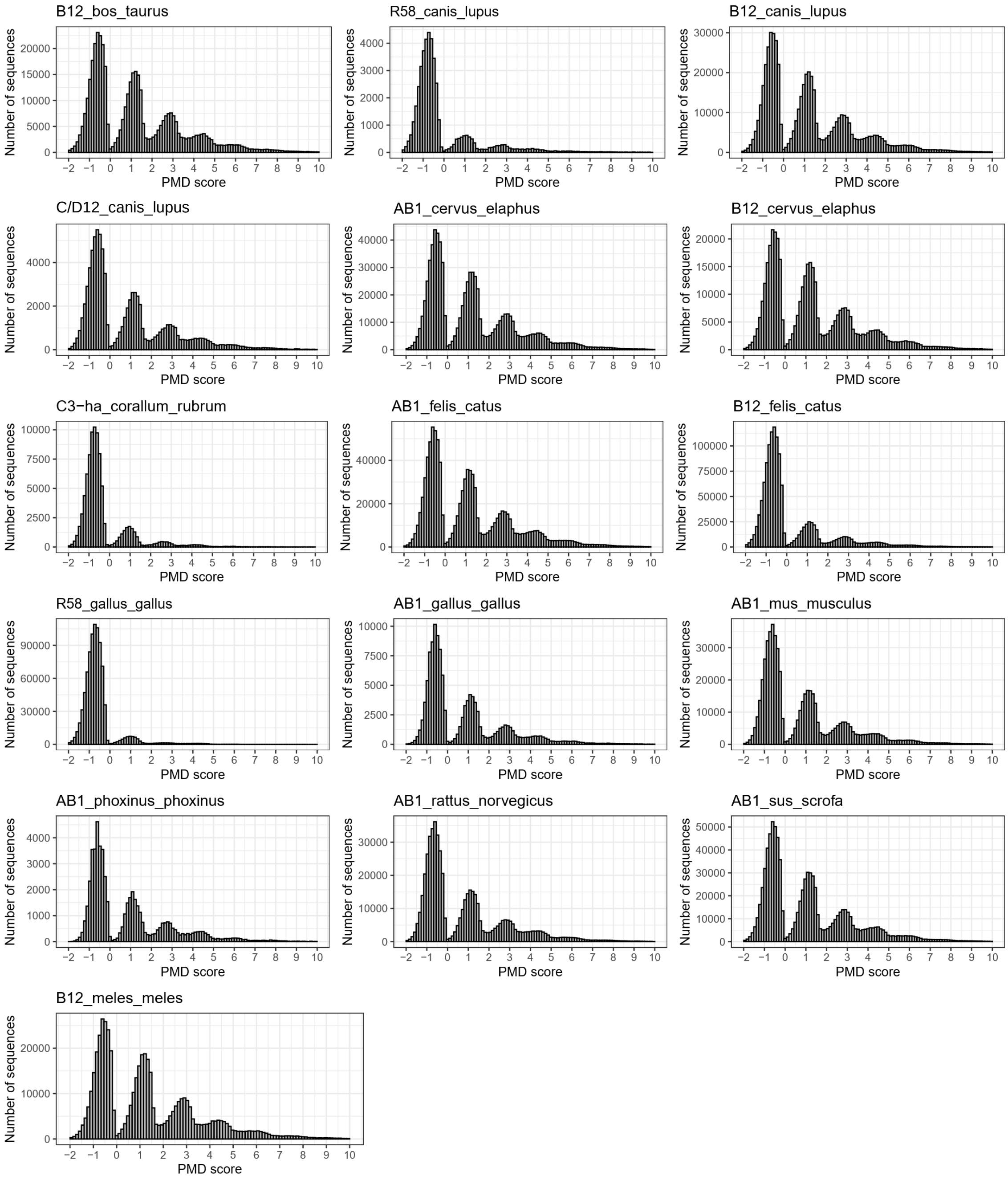


**Figure S6. PMDs distribution in the TS samples aligned to different animal species.** The distribution of the post-mortem damage (PMD) scores was computed only for those TS samples showing more than 100 contigs using the contig-BLAST approach (Dataset_S8).

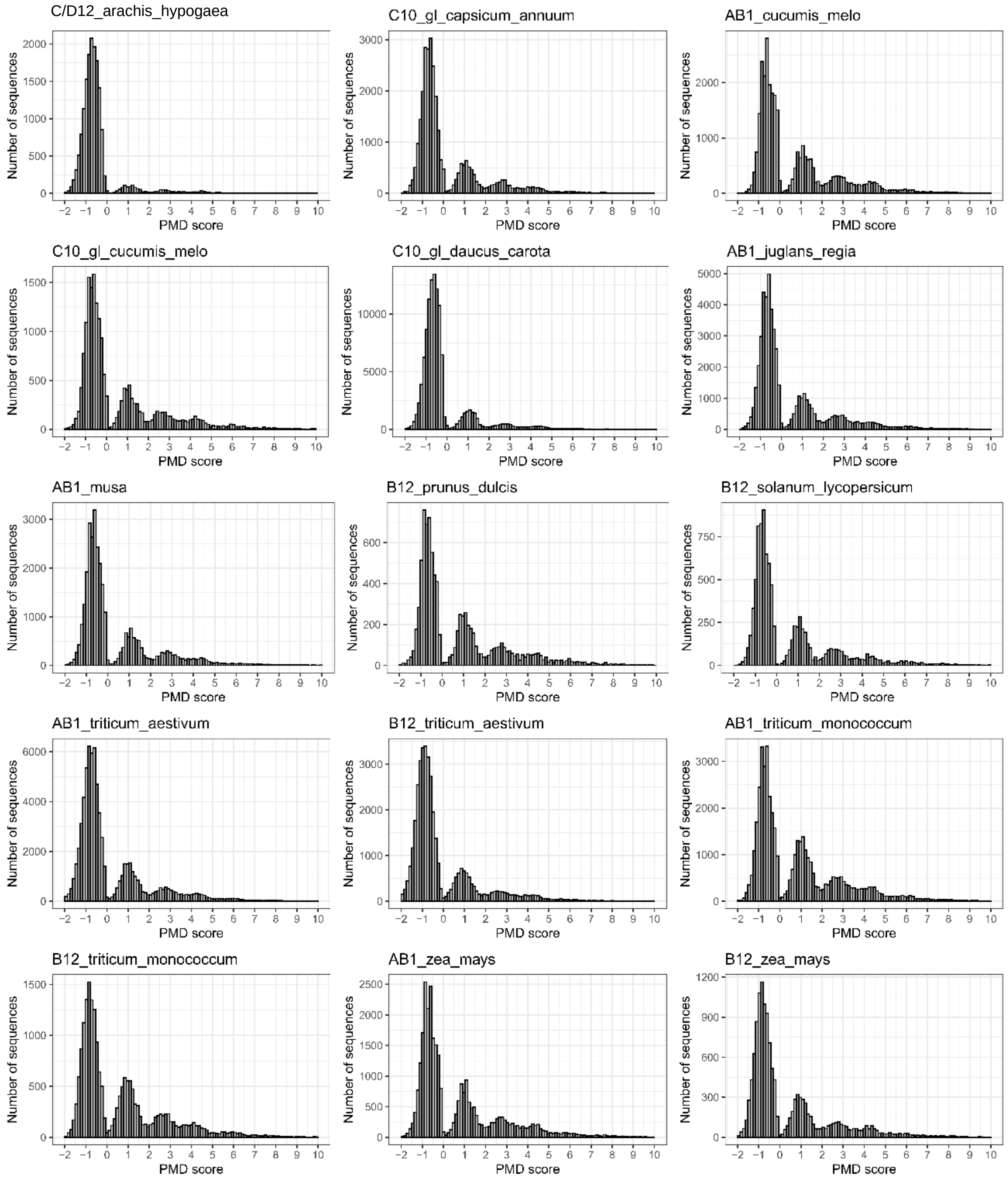


**Figure S7. PMDS distribution in the TS samples aligned to different plant species.** The distribution of the post-mortem damage (PMD) scores was computed only for those TS samples showing more than 100 contigs using the contig-BLAST approach (Dataset_S8).

### Tables

**Table S1. List of samples analyzed in this study (see Figure S1 for further details).**

| Sample ID[a] | Shroud section | Specimen | Genomic data | Human endogenous content (%) | Human average depth (X) | Meta genomics |
|---|---|---|---|---|---|---|
| AB1 | lateral edge | linen fragment | yes | 47.18 | 1.38 | yes |
| C/D12 | feet | linen fragment | yes | 5.64 | 0.10 | yes |
| A11 | feet | linen fragment | no | n.a. | n.a. | n.a. |
| C8 | arm | linen fragment | no | n.a. | n.a. | n.a. |
| C9 | arm | linen fragment | no | n.a. | n.a. | n.a. |
| C12 | feet | linen fragment | no | n.a. | n.a. | n.a. |
| R58 | Reliquary | linen fragments | yes | 0.57 | 0.02 | yes |
| C3_ha | hands | vacuum dust | yes | 1.93 | 0.05 | yes |
| C6_fa | face | empty | no | n.a. | n.a. | n.a. |
| C10_gl | glutei | vacuum dust | yes | 18.68 | 0.54 | yes |
| C12_fe | feet | vacuum dust | yes | 0.05 | 0.00 | yes |
| B12 | feet | microscope slide | yes | 39.60 | 1.14 | yes |

[a] The first code refers to the coordinates of the graphic map of the TS reported in the Figure S2 panel C of Barcaccia et al. [3]. The second code, if present, underlines the area of the body image from which the dust particles were vacuumed: hands (ha), face (fa), glutei (gl), feet (fe).

**Table S2. Kinship analyses.**

Genomic relatedness between the three best-performing TS samples (AB1, B12 and C10-gl) and the controls (Lab operator and collector).

| ind1 | ind2 | BREADR | READ2 | KIN (>0.05X) |
|---|---|---|---|---|
| AB1 | B12 | First_Degree | First_Degree | First_Degree |
| AB1 | collector | First_Degree | Same_Twins | First_Degree |
| AB1 | C10-gl | First_Degree | First_Degree | First_Degree |
| AB1 | operator | Unrelated | Unrelated | Unrelated |
| B12 | collector | First_Degree | First_Degree | First_Degree |
| B12 | C10-gl | First_Degree | First_Degree | First_Degree |
| B12 | operator | Unrelated | Unrelated | n.a. |
| C10-gl | collector | Same_Twins | Same_Twins | First_Degree |
| C10-gl | operator | Unrelated | Unrelated | Unrelated |

**Table S3. Summary of the Kraken2 results.**
Total number of analyzed reads and percentages of reads assigned to different domains for each sample.

| Sample | N. reads | Unassigned | Bacteria | Viruses | Archaea | Eukaryota | *Homo* |
|---|---|---|---|---|---|---|---|
| AB1 | 132 435 925 | 17.2 | 11.0 | 0.1 | 0.0 | 71.7 | 68.5 |
| C/D12 | 85 726 122 | 54.5 | 31.8 | 0.0 | 0.1 | 13.6 | 12.7 |
| R58 | 135 258 039 | 72.0 | 26.2 | 0.0 | 0.1 | 1.7 | 1.3 |
| C3_ha | 124 148 569 | 69.2 | 27.0 | 0.0 | 0.1 | 3.7 | 3.2 |
| C10_gl | 161 694 229 | 50.2 | 18.5 | 0.0 | 0.0 | 31.3 | 29.9 |
| C12_fe | 148 328 250 | 71.6 | 27.9 | 0.0 | 0.1 | 0.4 | 0.1 |
| B12 | 118 623 300 | 25.9 | 10.0 | 0.0 | 0.0 | 64.0 | 62.1 |

**Table S4. Summary metrics of the metagenomic assembly for each sample.**
The columns describe the total number of contigs obtained, the grand total of assembled base pairs, the N50 value of the assembly, the lengths of the longest and shortest contigs, and the average contig size.

| Sample | Total n. of contigs | Total bp | Min contig length (bp) | Max contig length (bp) | Avg contig length (bp) | N50 (bp) |
|---|---|---|---|---|---|---|
| AB1 | 4 711 086 | 1 143 646 932 | 118 | 68 555 | 242 | 240 |
| C/D12 | 284 933 | 80 382 933 | 146 | 155 175 | 282 | 240 |
| R58 | 348 254 | 80 015 682 | 151 | 254 563 | 229 | 194 |
| C3_ha | 353 365 | 89 037 482 | 151 | 254 543 | 251 | 213 |
| C10_gl | 955 934 | 196 561 234 | 150 | 254 691 | 205 | 185 |
| C12_fe | 298 238 | 73 208 206 | 150 | 254 505 | 245 | 214 |
| B12 | 4 693 106 | 1 169 249 982 | 118 | 271 010 | 249 | 249 |

**Table S5. Summary of the BLASTn analysis results.**

Total number of analyzed reads and the percentage of reads assigned to different domains for each sample.

| Sample | N. contigs | Unassigned | Bacteria | Viruses | Archea | Eukaryota | *Homo* |
|---|---|---|---|---|---|---|---|
| AB1 | 4 711 086 | 7.1 | 3.7 | 0.1 | 0.0 | 89.2 | 81.8 |
| C/D12 | 284 933 | 25.2 | 30.3 | 0.1 | 0.0 | 44.4 | 28.3 |
| R58 | 348 254 | 83.9 | 6.6 | 0.1 | 0.0 | 9.4 | 6.5 |
| C3_ha | 353 365 | 66.1 | 14.2 | 0.1 | 0.0 | 19.6 | 12.9 |
| C10_gl | 955 934 | 31.3 | 6.6 | 0.1 | 0.0 | 62.1 | 49.5 |
| C12_fe | 298 238 | 95.1 | 4.1 | 0.1 | 0.0 | 0.7 | 0.6 |
| B12 | 4 693 106 | 6.2 | 3.3 | 0.0 | 0.0 | 90.5 | 39.0 |

#### Datasets

**Dataset S1 (separate file).** Summary of genomic data (and 14C radiocarbon dating).

**Dataset S2 (separate file).** Summary of mitochondrial data.

**Dataset S3 (separate file).** Summary of proteomics analyses

**Dataset S4 (separate file).** Summary of metagenomic analyses (Archea, Bacteria, Fungi)

**Dataset S5 (separate file).** Outputs of Kraken.

**Dataset S6 (separate file).** Outputs of MetaPhlAn

**Dataset S7 (separate file).** Complete list of taxa identified using the contig-BLAST approach.

**Dataset S8 (separate file).** Summary of metagenomic analyses using the contig-BLAST approach (plant and animal taxa).
