## Supplementary material for "DNA Traces on the Shroud of Turin: Metagenomics of the 1978 Official Sample Collection": Dataset S5 and S6: Dataset_S5.html

Javascript must be enabled to view this page.

magnitude
magnitudeUnassigned

AB1 metaphlan results
C3 metaphlan results
A11 metaphlan results
C3\_ha metaphlan results
C10\_gl metaphlan results
C12\_fe metaphlan results
B12 metaphlan results

289316044842326098319320777678947905030710616245725200670

126958543963405197228318739918747645948410555178315825787

0000000
1432078384753861030836372029525908236106748825403

23883204732246825590
0000

0000
23883204732246825590

0000
23883204732246825590

0000
23883204732246825590

0000
23883204732246825590

0000
23883204732246825590

23883204732246825590

15132
0

15132
0

0
15132

15132
0

0
15132

0
15132

15132

0000000
31008110956191030889583710852985721959288

4028791627065657472743772151358145749
0000000

4028791627065657472743772151358145749
0000000

12030524506565724427256915135827441
0000000

00
1203017097

12030
0

12030

17097
0

17097

10344
0

10344
0

10344

5245065657244272569151358
00000

0
52450

52450

2217018308
00

2217018308

34607177521915823644
0000

34607177521915823644

0000
8880667565339406

8880667565339406

00000
282578638202284712030118308

0000
24834228471203061550

0000
24834228471203061550

24834228471203061550

00
7821525101

62107
0

62107

0
25101

25101

16108
0

16108

1222314817
00

00
1222314817

1222314817

16034
0

0
16034

16034

760771
0

0
33098

33098

727673
0

727673

0
16840

16840
0

16840

000000
60551927624113141501455719393

000000
60551927624113141501455719393

60551927624113141501455719393
000000

000000
60551927624113141501455719393

000000
60551927624113141501455719393

60551927624113141501455719393

00
9387113672

9387113672
00

00
9387113672

00
9387113672

113672
0

113672

9387
0

9387

14433133181126412047149709471
000000

14433133181126412047149709471
000000

000000
14433133181126412047149709471

000000
14433133181126412047149709471

000000
14433133181126412047149709471

14433133181126412047149709471

0
7844

7844
0

7844
0

0
7844

7844
0

7844

16172
0

16172
0

0
16172

0
16172

16172
0

16172

00000
254352319682314928032769411

0
4956

0
4956

4956
0

0
4956

4956

00000
254352319682314928032764455

00
1533436666

22486
0

22486
0

22486

00
1533414180

14180
0

14180

0
15334

15334

1396861745
00

0
48014

0
19109

19109

28905
0

28905

00
1396813731

13968
0

13968

13731
0

13731

12643
0

12643
0

12643
0

12643

3196891818312
000

0
31968

31968
0

31968

91818312
00

91818312
00

91818312

17288328032291997
000

00
2498480478

11911
0

11911

37292
0

37292

0
28305

28305

0
13073

13073

0
14881

14881

000
1048432803231110

0
8645

8645

1289716828
00

1289716828

00
1434614282

1434614282

0
24071

24071

10705
0

10705

16114
0

16114

00
1806510816

1806510816

17216
0

17216

00
881146001

00
88116494

88116494

0
39507

39507

00
952324184

00
95239416

95239416

0
14768

14768

0
6894

0
6894

6894

00
2472285293

976942380
00

976942380

1495342913
00

1495342913

0
18037

0
18037

18037

00
66135353092

11560
0

11560
0

11560

0
11271

11271
0

11271

66135330261
00

0
8667

8667

0
64557

64557

0
28834

28834

0
15181

15181

10386
0

10386

45662
0

45662

0
13598

13598

754411195
00

754411195

29627
0

29627

2265847450
00

2265847450

0
8869

8869

10545
0

10545

0
43516

43516

12085
0

12085

16022
0

16022

12434
0

12434
0

12434
0

0
12434

0
12434

12434

0
14379

14379
0

14379
0

0
14379

0
14379

14379

0
14505

0
14505

14505
0

0
14505

14505
0

0
14505

14505

16443535107326398804289501711914472985891423601
0000000

1385949258976075495151272524202815149405589
0000000

745111648
00

00
745111648

7451
0

0
7451

7451

0
11648

0
11648

11648

27472175442
00

00
27472175442

00
27472175442

0
27472

27472

175442
0

175442

896377226630638421464293121444162505
000000

000000
896377226630638421464293121444162505

193612032450742
000

0
17280

17280

000
193612032433462

193612032433462

5689215454
00

0
15483

15483

0
15090

15090

13588
0

13588

00
1273115454

1273115454

21254
0

21254
0

21254

82012422663063842144396912144475055
000000

37830
0

37830

56877674921
00

56877674921

0
16163

16163

1194403734633142144396912144440188
000000

1194403734633142144396912144440188

0000
774431143633070018704

774431143633070018704

16635
0

16635

000
195171530015150

195171530015150
000

00
1277615300

1277615300
00

1277615300

0
6741

0
6741

6741

0
15150

0
15150

15150

226121073311165310919702301514974935
0000000

0000000
226121073311165310919702301514974935

11648
0

11648
0

11648

00
5881447025

23699
0

23699

23326
0

23326

58814
0

58814

11736116531091911416151497969
000000

000000
11736116531091911416151497969

11736116531091911416151497969

2261295595
00

00
2261239948

2261239948

55647
0

55647

0
8293

8293
0

8293

134733776062709
000

00
3776036529

36529
0

0
36529

36529

0
37760

37760
0

37760

26180
0

0
26180

26180
0

26180

0
13473

0
13473

13473
0

13473

000
5908951145023585

22554511450
00

22554511450
00

00
22554511450

22554511450

16572
0

16572
0

0
16572

16572

1996323585
00

00
1996323585

23585
0

23585

19963
0

19963

3263471531147300623505445923
00000

00000
3196521531147300623505445923

0
83423

83423
0

83423

0
19504

0
19504

19504

00000
3001481407951300623505440204

33230
0

33230

00
448121335079

448121335079

0
72872

72872

40204
0

40204

28934
0

28934

107384
0

107384

30062
0

30062

0
35054

35054

0
30942

30942

0
54846

34336

20510

5719
0

5719
0

5719

39773
0

0
39773

39773

6695
0

0
6695

6695
0

6695

0
9134

9134
0

9134
0

0
9134

9134

21062
0

21062
0

21062
0

0
21062

21062

509291418762727
000

509291418762727
000

000
509291418762727

000
509291418762727

000
509291418762727

509291418762727

0000000
15472175704229250124183288357212598467314

6586
0

0
6586

6586
0

6586
0

6586

782457527
00

00
782457527

0
26914

26914
0

26914

00
782430613

0
7824

7824

30613
0

30613

321183687928357794137886311985
000000

000000
321183687928357794137886311985

000
96721945911985

967219459
00

967219459

0
11985

11985

0
34768

34768
0

34768

00000
3211836879283573497359404

20414786917503
000

20414786917503

0000
36879283572710441901

36879283572710441901

0
11704

11704

1547214358619237189240173431133735397802
0000000

0
14134

14134
0

0
14134

14134

95053601129772
000

95053601129772
000

95053601129772
000

95053601129772

0
5967

5967
0

5967
0

5967

000000
3518084005409326113274394101599

000000
3518084005409326113274394101599

229911730422212
000

229911730422212

351802921117433212333598679387
000000

351802921117433212333598679387

31803234992259538408
0000

31803234992259538408

85095
0

0
43808

0
43808

43808

41287
0

0
41287

41287

000000
723951083664830811229959341167202

72395442971511966555167202
00000

7239533159142434
000

7239533159142434

2103117697
00

2103117697

0
24768

24768

232661511915699
000

232661511915699

64069331894574459341
0000

0000
64069331894574459341

64069331894574459341

27689
0

27689
0

27689
0

10329
0

10329

0
17360

17360

0000000
183123232767580573125810965267061889456163

00
2783442278

27834
0

27834
0

27834
0

27834

0
42278

0
42278

0
42278

42278

00000
5178342257299985558420865

13697
0

0
13697

13697
0

13697

517834225729998555847168
00000

0000
51783422572999842808

10751
0

10751

00
1365621762

1365621762

0000
14247204951701615936

14247204951701615936

000
131291298213842

131291298213842

0
13030

13030

7168
0

0
7168

7168

0
12776

12776
0

12776

0000000
103506163746270373122316758976751587416456

86547901395832430861405974037918715
0000000

00000
5832430861277584037918715

00000
4501630861277584037918715

4501630861277584037918715

13308
0

13308

8654
0

8654
0

8654

12839
0

12839
0

12839

0
790139

0
790139

790139

00000
68492548152106116533386760434

137320212617312280
000

89931212617312280
000

89931212617312280

0
47389

47389

0
88174

0
88174

88174

000
21301212683848548

0
49013

49013

00
21301116057

21301116057

0
848548

848548

47613
0

47613

0
17901

17901
0

17901

223091099752158742533
0000

22309872332158742533
0000

22309872332158742533

0
22742

22742

0
24882

24882
0

24882

88203324612049928222015
00000

0000
882012049928210839

0000
882012049928210839

882012049928210839

3324611176
00

00
3324611176

3324611176

000
1715211858740288

1715240288
00

00
1715240288

1715240288

118587
0

0
80247

80247

0
38340

38340

000000
175402659251047079671611208275004

0
11208

11208
0

11208

0
47065

47065
0

47065

00
134858168850

0
62955

62955

0
29319

29319

71903139531
00

71903139531

17540
0

17540
0

17540

84388626775191159089
0000

84388626775191159089
0000

59089

843886267751911

00
4203044805

00
4203044805

4203044805

0
46679

46679
0

46679

60567810200494473191030218842
000000

00
60567811537

605678
0

605678
0

605678

11537
0

11537
0

11537

1020049447319103027305
00000

00000
1020049447319103027305

00000
1020049447319103027305

1020049447319103027305

0
8880

0
8880

0
8880

0
8880

0
8880

8880

1348683928953
000

000
1348683928953

1348683928953
000

1348683928953
000

1348683928953
000

1348683928953

0
31808

0
31808

0
31808

31808
0

0
31808

31808

375754028840
000

0
3757

0
3757

0
3757

3757
0

0
3757

3757

5357
0

5357
0

0
5357

0
5357

5357
0

5357

0
5402

0
5402

5402
0

5402
0

5402
0

5402

0
3483

3483
0

0
3483

3483
0

3483
0

3483

850098116506873793796021786943142007744644945
0000000

0
5144

0
5144

0
5144

0
5144

5144
0

5144

0000000
753761674063530826072177912227430730

753761674063530826072177912227430730
0000000

753761674063530826072177912227430730
0000000

0000000
753761674063530826072177912227430730

753761674063530826072177912227430730
0000000

753761674063530826072177912227430730

0000000
841448114832813209695612366593761490204557171

00000
2262611775605941285563209

00000
2262611775605941285563209

447571285553910
000

0
26668

26668

4475712855
00

4475712855

15389
0

15389

11853
0

11853

0
9299

9299
0

9299

000
226261177515837

00
67627329

67627329

5730
0

5730

000
10134117758508

10134117758508

000
26453016403253031

26453016403253031
000

27866
0

0
27866

27866

18602
0

18602
0

18602

1160735087
00

0
11607

11607

15112
0

15112

0
11171

11171

0
8804

8804

12872
0

0
12872

12872

0
9303

9303
0

9303

38762
0

0
38762

38762

0
9367

0
9367

9367

14566
0

14566
0

14566

13729
0

13729
0

13729

00
866684030

0
25599

25599

00
866658431

866658431

1963711640316028
000

2608516403
00

2608516403

30303
0

30303

19568
0

19568

5838116028
00

5090416028

7477

9693
0

9693

15968
0

15968

18393
0

18393

17980
0

17980

00
119188787

00
119188787

11918

8787

63352211122506183091168990156554539861821873
0000000

0000000
63352211122506183091168990156554539861821873

0
96133

0
96133

96133

11676
0

0
11676

11676

0
15182

15182
0

15182

14281
0

0
14281

14281

0000000
63352211122506183091153808156554539861699783

342231167015764
000

342231167015764

00000
11229836907159001323243121

11229836907159001323243121

48960511762619393149711578339852
000000

48960511762619393149711578339852

5563564921644163698122937127539539861566226
0000000

5563564921644163698122937127539539861566226

554553465934820
000

554553465934820

0
80076

80076

0000000
250379741338199615861926629554643358330

1137754809068021123289231654140783
000000

0
29605

29605
0

29605

000000
1520325201345009694715929721214

0
118058

118058

1520325201345009694721214
00000

1520325201345009694721214

16499
0

16499

24740
0

24740

000000
399302288933521158631606853839

23972228891630833420
0000

23972228891630833420

1595820419
00

1595820419

0
17213

17213

1586316068
00

1586316068

0000
58642104795628914580

00
1386313124

1386313124

202201654614580
000

202201654614580

0
15889

15889

245591047910730
000

9262

152971047910730

21545
0

0
21545

21545

39905260431397535330346415464379448
0000000

0
13836

13836
0

13836

0000
13975133741447221329

13975133741447221329
0000

13975133741447221329

16880
0

16880
0

16880

11538
0

0
11538

11538

6788
0

0
6788

6788

162372604354074
000

19421
0

19421

000
162372604334653

162372604334653

219562016933314
000

000
219562016933314

219562016933314

7460
0

7460
0

7460
0

7460

00
7453588160

7453588160
00

00
1181023982

1181023982

22403
0

22403

14747
0

14747

26371
0

26371

0
13483

13483

2557524324
00

2557524324

00
147049013

14704
0

0
14704

14704

0
9013

0
9013

9013

40926
0

0
40926

0
21466

21466

19460
0

19460

00
60885758

00
60885758

00
60885758

00
60885758

60885758

13062
0

0
13062

13062
0

13062
0

13062

00
1327516656

1327516656
00

00
1327516656

00
1327516656

1327516656

152845028664238019159971207269346332044072
0000000

20737331227171841353078000120628
000000

000000
1944723122717184135307800069164

000000
1944723122717184135307800069164

1944723122717184135307800069164

00
1290151464

1290112811
00

1290112811

0
38653

38653

17698717493463319954
0000

0
71749

0
71749

71749

0
34633

0
34633

34633

00
1769819954

1769819954
00

1769819954

000
2614738700160341

000
2614738700160341

9918
0

9918

79652
0

79652

261473870034834
000

261473870034834

35937
0

35937

000000
127723221671520835746921292691743149

000000
126133021671520835662551292691743149

0
11220

11220

000
762151722364825

762151722364825

38234
0

38234

1114429085
00

1114429085

0
11363

11363

0
16755

16755

0
17315

17315

11772125583
00

11772125583

0
16268

16268

15238
0

15238

00
1242232749

1242232749

33845
0

33845

7747
0

7747

00
2019231051

2019231051

1172752511518870
000

1172752511518870

0
11627

11627

0
6830

6830

000000
27311660517208351645313577121806

2333018333

24978642184208351645313577121806

12643
0

12643

11006
0

11006

19827
0

19827

8208
0

8208

130524383221276224448636004
00000

130524383221276224448636004

0
14372

14372

13499
0

13499

0
13287

13287

00
1708556401

1708556401

8000
0

8000

25833
0

25833

0
35597

7616

27981

0
11617

11617

33949195781372717599
0000

33949195781372717599

00
10319279527

10319279527

0
12348

12348

18285
0

18285

0
10974

10974

15645
0

15645

0
15859

15859

0000
86848268411489818487

86848268411489818487

17683
0

17683

0
10296

10296

000
11729229119135747

7022429119

47068135747

22479
0

22479

5202130171152138500
0000

5202130171152138500

66111
0

66111

0
8437

0
8437

8437

5150
0

0
5150

5150

10752
0

10752
0

10752

0
7103

0
7103

7103
0

0
7103

0
7103

7103

00
1112418268

00
1112418268

111246236
00

0
11124

0
11124

11124

0
6236

6236
0

6236

12032
0

0
12032

0
12032

12032

00000
2310214870171472948026529

00000
2310214870171472948026529

2310214870171472948026529
00000

2310214870171472948026529
00000

2310214870171472948026529
00000

2310214870171472948026529

2257138279
00

00
2257138279

00
2257138279

9985
0

9985
0

0
9985

9985

2257118247
00

2257118247
00

1099118247
00

1099118247

11580
0

11580

0
10047

10047
0

10047
0

10047

0
17157

17157
0

0
17157

17157
0

0
17157

17157
0

17157

0
15734

0
15734

0
15734

15734
0

15734
0

15734
0

15734

8827
0

8827
0

0
8827

0
8827

8827
0

8827
0

8827

0
2664

2664
0

0
2664

0
2664

2664
0

0
2664

2664

0
26562

18133
0

18133
0

0
18133

0
18133

0
18133

18133

0
8429

0
8429

0
8429

0
8429

0
8429

8429

000000
37984186217541256821067905598361644791

10196
0

10196
0

0
10196

10196
0

10196
0

10196

0
5379

0
5379

5379
0

0
5379

0
5379

5379

0
8702

0
8702

8702
0

8702
0

0
8702

8702

0
14457

0
14457

14457
0

3494
0

3494
0

3494

0
10963

0
10963

10963

5897
0

5897
0

5897
0

5897
0

5897
0

5897

0
17214

17214
0

0
17214

0
17214

0
17214

17214

0
5969

5969
0

0
5969

0
5969

0
5969

5969

000000
29780954139378990596885548118813517

000000
26165143969746975934707391557297252

00
215397547

215397547
00

61237547
00

61237547

7900
0

7900

0
7516

7516

57386256297
00

0
57386

57386
0

57386

0
256297

0
232451

208243

24208

23846
0

23846

2568518327567697593470796091297252
000000

123249499
00

00
123249499

123249499

000000
2556194327567697593470796091287753

24341
0

24341

709903963125057
000

709903963125057

00
3626012277

3626012277

2667526499415771149181442290475
000000

2667526499415771149181442290475

0
15776

15776

00
15790229529

15790229529

17144
0

17144

14023
0

14023

412541699214426
000

412541699214426

124006310313110467
000

124006310313110467

0000
110584226441485031613

110584226441485031613

0
32754

32754

1843014346
00

11263

716714346

0
27707

27707

0
13784

13784

0
20767

20767

4996996763822146197892059698129
000000

4996996763822146197892059698129

0
14424

14424
0

14424
0

14424

0
12033

0
12033

0
12033

12033

00
1202131622

31622
0

31622
0

31622

12021
0

12021
0

12021

361581169632014662178156561516265
000000

000000
22571616963201462473059461165388

21184916963201462473059461151948
000000

13079
0

13079

00
87868883

87868883

0000
4005720146936226496

4005720146936226496

8632
0

8632

0
8727

8727

00
991410762

991410762

1114913768
00

1114913768

1437991946720
000

1437991946720

0
12453

12453

212248588
00

212248588

0
10392

10392

173061536813587
000

173061536813587

22009169631018412257
0000

22009169631018412257

2275969790
00

2275969790

12163
0

12163

1386713440
00

1386713440
00

1386713440

00
46880191285

0
19980

0
19980

19980

00
29928103815

42311
0

42311

0
8630

8630

00
731510204

731510204

0
24119

24119

7169
0

7169

1398320012
00

1398320012

5785
0

0
5785

5785

1003412985
00

00
1003412985

1003412985

0
6918

0
6918

6918

48720
0

0
11863

11863

18796
0

18796

18061
0

18061

3414885014
00

0
6512

0
6512

6512

00
1098956240

1098914241
00

1098914241

0
20277

20277

11118
0

11118

0
10604

10604

00
2315922262

00
972411960

972411960

00
1343510302

1343510302

3717897798193041664
0000

0
81930

81930
0

81930

1915741664
00

1915741664
00

1915741664

180219779
00

00
82239779

82239779

9798
0

9798

0000
17659276691517032914

0000
17659276691517032914

276691517012706
000

276691517012706

1765920208
00

1765920208

0
12603

0
12603

0
12603

12603
0

0
12603

12603

000
5546654509173908

3308788555
00

3308788555
00

0
15352

0
15352

15352

18394
0

18394
0

18394

16247
0

0
16247

16247

0
9500

9500
0

9500

0
21877

21877
0

21877

32037
0

17008
0

17008

15029
0

15029

8235
0

0
8235

8235

000
74375450978062

00
5450917357

00
5450917357

0
17357

17357

54509
0

54509

0
25427

17644
0

0
17644

17644

0
7783

7783
0

7783

35278
0

0
23940

0
23940

23940

0
11338

11338
0

11338

7437
0

7437
0

7437
0

7437

149427291
00

00
81617291

8161
0

8161
0

8161

7291
0

0
7291

7291

0
6781

0
6781

0
6781

6781

0000
109449334151171871905

1000021171871905
000

000
1000021171871905

1835911450
00

11450
0

11450

18359
0

18359

000
719141171811785

11945
0

11945

12028
0

12028

0
6201

6201

10167
0

10167

000
315731171811785

315731171811785

33072
0

33072
0

33072

972915598
00

00
972915598

972915598

944733415
00

0
33415

33415
0

33415
0

33415

0
9447

9447
0

9447
0

9447

00000
655408119893357779905505044

00000
641174119893357779905473196

00000
641174119893357779905473196

7607
0

0
7607

7607

714082195139773
000

000
714082195139773

15454

559542195139773

00
2711240741

66028519
00

66028519

14713
0

14713

10477
0

10477

1003317509
00

1003317509

0
8498

8498
0

8498

00000
42379359070206919905257557

51417
0

51417

0
5997

5997

169439342
00

169439342

00000
125563354832069199057831

125563354832069199057831

1852015073
00

1852015073

1140312294
00

1140312294

0
7903

7903

5025489709562
000

5025489709562

9229
0

9229

3231335727
00

1448712218

1782623509

1200331877
00

1200331877

720581461739042
000

720581461739042

00
2221621632

2221621632

3484217763
00

3484217763

10546
0

10546

0
21605

15083
0

15083

0
6522

6522

0000
1112543887215086105022

82864388721508618593
0000

82864388721508618593

818912455
00

818912455

4983
0

4983

0
12413

12413

16450
0

16450

0
15332

15332

2020124796
00

2020124796

00
1423431848

1423431848
00

00
451719328

00
451719328

451719328

971712520
00

9717
0

9717

12520
0

12520

00
4062213905

4062213905
00

00
4062213905

1663413905
00

1663413905
00

0
11941

11941

4693
0

4693

0
13905

13905

0
23988

23988
0

10222
0

10222

0
13766

13766

19149673138236016655725549480
00000

6696511060116016655725549480
00000

6696511060116016655725549480
00000

6696511060116016655725549480
00000

6696511060116016655725549480
00000

6696511060116016655725549480
00000

4683741060116016655725458251
00000

4683741060116016655725458251

00
20127791229

20127791229

1245316207812
00

00
1245316207812

1245316207812
00

00
1245316207812

00
1245316207812

1245316207812
00

1245316207812
