## Supplementary material for "DNA Traces on the Shroud of Turin: Metagenomics of the 1978 Official Sample Collection": Dataset S5 and S6: Dataset_S6.html

Javascript must be enabled to view this page.

magnitude
magnitudeUnassigned

AB1 metaphlan results
C/D12 metaphlan results
R58 metaphlan results
C3\_ha metaphlan results
C10\_gl metaphlan results
C12\_fe metaphlan results
B12 metaphlan results

289316044842326098319320777678947905030710616245725200670

126958543963405197228318739918747645948410555178315825787

00000
19149673138236016655725549480

00000
6696511060116016655725549480

6696511060116016655725549480
00000

00000
6696511060116016655725549480

00000
6696511060116016655725549480

00000
6696511060116016655725549480

4683741060116016655725458251
00000

4683741060116016655725458251

00
20127791229

20127791229

1245316207812
00

1245316207812
00

00
1245316207812

00
1245316207812

00
1245316207812

1245316207812
00

1245316207812

1432078384753861030836372029525908236106748825403
0000000

17157
0

17157
0

17157
0

0
17157

0
17157

17157
0

17157

0
14505

14505
0

14505
0

14505
0

14505
0

0
14505

14505

2664
0

2664
0

2664
0

2664
0

0
2664

0
2664

2664

0000000
850098116506873793796021786943142007744644945

1112418268
00

1112418268
00

111246236
00

0
6236

0
6236

6236

0
11124

0
11124

11124

0
12032

0
12032

0
12032

12032

0000000
753761674063530826072177912227430730

0000000
753761674063530826072177912227430730

0000000
753761674063530826072177912227430730

0000000
753761674063530826072177912227430730

753761674063530826072177912227430730
0000000

753761674063530826072177912227430730

0
7103

7103
0

0
7103

7103
0

0
7103

7103

2310214870171472948026529
00000

2310214870171472948026529
00000

00000
2310214870171472948026529

00000
2310214870171472948026529

2310214870171472948026529
00000

2310214870171472948026529

0
5144

5144
0

0
5144

0
5144

0
5144

5144

841448114832813209695612366593761490204557171
0000000

26453016403253031
000

000
26453016403253031

000
1963711640316028

0
18393

18393

17980
0

17980

0
19568

19568

2608516403
00

2608516403

0
15968

15968

30303
0

30303

5838116028
00

5090416028

7477

0
9693

9693

9303
0

0
9303

9303

00
119188787

119188787
00

8787

11918

13729
0

13729
0

13729

27866
0

27866
0

27866

18602
0

18602
0

18602

00
1160735087

11171
0

11171

15112
0

15112

0
8804

8804

0
11607

11607

0
12872

0
12872

12872

0
38762

38762
0

38762

0
9367

9367
0

9367

00
866684030

00
866658431

866658431

0
25599

25599

0
14566

14566
0

14566

0
13062

0
13062

13062
0

0
13062

13062

0000000
152845028664238019159971207269346332044072

17698717493463319954
0000

34633
0

0
34633

34633

1769819954
00

1769819954
00

1769819954

0
71749

0
71749

71749

000000
20737331227171841353078000120628

1944723122717184135307800069164
000000

000000
1944723122717184135307800069164

1944723122717184135307800069164

00
1290151464

0
38653

38653

00
1290112811

1290112811

000000
127723221671520835746921292691743149

10752
0

0
10752

10752

000000
126133021671520835662551292691743149

0
22479

22479

27311660517208351645313577121806
000000

24978642184208351645313577121806

2333018333

18285
0

18285

0
12643

12643

0
33845

33845

14372
0

14372

19827
0

19827

0
66111

66111

0
15859

15859

0
7747

7747

00
2019231051

2019231051

11627
0

11627

0
12348

12348

10974
0

10974

0000
33949195781372717599

33949195781372717599

0
25833

25833

000
762151722364825

762151722364825

00000
130524383221276224448636004

130524383221276224448636004

86848268411489818487
0000

86848268411489818487

1242232749
00

1242232749

0
17315

17315

00
11772125583

11772125583

0
13287

13287

0
15238

15238

6830
0

6830

1172752511518870
000

1172752511518870

00
1114429085

1114429085

0
11617

11617

0
8000

8000

35597
0

7616

27981

0
13499

13499

0
11363

11363

0
11006

11006

0
17683

17683

5202130171152138500
0000

5202130171152138500

0
10296

10296

0
11220

11220

10319279527
00

10319279527

0
8208

8208

0
16268

16268

000
11729229119135747

7022429119

47068135747

16755
0

16755

0
15645

15645

38234
0

38234

00
1708556401

1708556401

5150
0

5150
0

5150

0
8437

0
8437

8437

2614738700160341
000

2614738700160341
000

79652
0

79652

000
261473870034834

261473870034834

9918
0

9918

0
35937

35937

2262611775605941285563209
00000

2262611775605941285563209
00000

000
226261177515837

67627329
00

67627329

10134117758508
000

10134117758508

0
5730

5730

447571285553910
000

15389
0

15389

0
26668

26668

0
11853

11853

00
4475712855

4475712855

0
9299

0
9299

9299

250379741338199615861926629554643358330
0000000

00
7453588160

7453588160
00

0
14747

14747

0
13483

13483

00
1181023982

1181023982

00
2557524324

2557524324

26371
0

26371

22403
0

22403

000000
1137754809068021123289231654140783

21545
0

21545
0

21545

58642104795628914580
0000

000
202201654614580

202201654614580

000
245591047910730

9262

152971047910730

0
15889

15889

1386313124
00

1386313124

000000
399302288933521158631606853839

00
1595820419

1595820419

23972228891630833420
0000

23972228891630833420

1586316068
00

1586316068

0
17213

17213

000000
1520325201345009694715929721214

0
24740

24740

118058
0

118058

00000
1520325201345009694721214

1520325201345009694721214

16499
0

16499

0
29605

0
29605

29605

0
40926

40926
0

21466
0

21466

0
19460

19460

7460
0

0
7460

0
7460

7460

39905260431397535330346415464379448
0000000

0
11538

11538
0

11538

000
219562016933314

219562016933314
000

219562016933314

0000
13975133741447221329

13975133741447221329
0000

13975133741447221329

6788
0

6788
0

6788

13836
0

0
13836

13836

0
16880

16880
0

16880

000
162372604354074

0
19421

19421

162372604334653
000

162372604334653

00
147049013

9013
0

0
9013

9013

14704
0

0
14704

14704

1327516656
00

1327516656
00

1327516656
00

00
1327516656

1327516656

60885758
00

60885758
00

60885758
00

60885758
00

60885758

63352211122506183091168990156554539861821873
0000000

63352211122506183091168990156554539861821873
0000000

0000000
63352211122506183091153808156554539861699783

342231167015764
000

342231167015764

000
554553465934820

554553465934820

5563564921644163698122937127539539861566226
0000000

5563564921644163698122937127539539861566226

11229836907159001323243121
00000

11229836907159001323243121

48960511762619393149711578339852
000000

48960511762619393149711578339852

80076
0

80076

0
14281

0
14281

14281

15182
0

0
15182

15182

11676
0

11676
0

11676

96133
0

96133
0

96133

0
26562

0
8429

0
8429

0
8429

8429
0

8429
0

8429

18133
0

18133
0

18133
0

0
18133

0
18133

18133

31008110956191030889583710852985721959288
0000000

16172
0

0
16172

0
16172

0
16172

16172
0

16172

0
12434

0
12434

12434
0

12434
0

0
12434

12434

9387113672
00

9387113672
00

00
9387113672

9387113672
00

0
9387

9387

0
113672

113672

000000
14433133181126412047149709471

14433133181126412047149709471
000000

14433133181126412047149709471
000000

000000
14433133181126412047149709471

000000
14433133181126412047149709471

14433133181126412047149709471

7844
0

0
7844

7844
0

7844
0

0
7844

7844

00000
254352319682314928032769411

254352319682314928032764455
00000

12643
0

12643
0

0
12643

12643

17288328032291997
000

881146001
00

0
39507

39507

88116494
00

88116494

000
1048432803231110

00
1289716828

1289716828

0
17216

17216

00
1434614282

1434614282

0
8645

8645

16114
0

16114

24071
0

24071

0
10705

10705

00
1806510816

1806510816

00
2472285293

00
1495342913

1495342913

00
976942380

976942380

00
952324184

95239416
00

95239416

14768
0

14768

6894
0

6894
0

6894

0
18037

0
18037

18037

00
2498480478

13073
0

13073

0
11911

11911

0
28305

28305

14881
0

14881

37292
0

37292

000
3196891818312

00
91818312

91818312
00

91818312

31968
0

31968
0

31968

1396861745
00

48014
0

28905
0

28905

19109
0

19109

00
1396813731

0
13731

13731

13968
0

13968

00
66135353092

11271
0

11271
0

11271

00
66135330261

0
28834

28834

0
45662

45662

0
43516

43516

10545
0

10545

0
29627

29627

0
12085

12085

64557
0

64557

8667
0

8667

0
16022

16022

00
2265847450

2265847450

8869
0

8869

754411195
00

754411195

0
10386

10386

13598
0

13598

0
15181

15181

11560
0

11560
0

11560

00
1533436666

0
22486

0
22486

22486

00
1533414180

0
15334

15334

0
14180

14180

4956
0

0
4956

0
4956

0
4956

4956

4028791627065657472743772151358145749
0000000

4028791627065657472743772151358145749
0000000

12030524506565724427256915135827441
0000000

5245065657244272569151358
00000

52450
0

52450

34607177521915823644
0000

34607177521915823644

2217018308
00

2217018308

0000
8880667565339406

8880667565339406

1203017097
00

17097
0

17097

0
12030

12030

0
10344

0
10344

10344

00000
282578638202284712030118308

1222314817
00

1222314817
00

1222314817

7821525101
00

0
25101

25101

16108
0

16108

0
62107

62107

0
16034

16034
0

16034

16840
0

0
16840

16840

760771
0

0
727673

727673

33098
0

33098

0000
24834228471203061550

0000
24834228471203061550

24834228471203061550

14379
0

14379
0

0
14379

0
14379

0
14379

14379

60551927624113141501455719393
000000

000000
60551927624113141501455719393

000000
60551927624113141501455719393

60551927624113141501455719393
000000

60551927624113141501455719393
000000

60551927624113141501455719393

0000
23883204732246825590

0000
23883204732246825590

23883204732246825590
0000

23883204732246825590
0000

23883204732246825590
0000

23883204732246825590
0000

23883204732246825590

15132
0

15132
0

0
15132

15132
0

0
15132

15132
0

15132

4062213905
00

00
4062213905

00
4062213905

00
1663413905

1663413905
00

0
11941

11941

0
13905

13905

4693
0

4693

0
23988

0
23988

0
13766

13766

10222
0

10222

37984186217541256821067905598361644791
000000

5546654509173908
000

000
74375450978062

00
5450917357

00
5450917357

54509
0

54509

17357
0

17357

35278
0

23940
0

23940
0

23940

11338
0

0
11338

11338

0
25427

7783
0

0
7783

7783

0
17644

17644
0

17644

0
7437

7437
0

0
7437

7437

00
3308788555

3308788555
00

0
18394

0
18394

18394

16247
0

0
16247

16247

15352
0

15352
0

15352

0
9500

0
9500

9500

21877
0

21877
0

21877

0
8235

0
8235

8235

32037
0

15029
0

15029

0
17008

17008

00
149427291

6781
0

0
6781

6781
0

6781

81617291
00

0
7291

0
7291

7291

8161
0

0
8161

8161

00000
655408119893357779905505044

00000
641174119893357779905473196

00000
641174119893357779905473196

000
714082195139773

000
714082195139773

15454

559542195139773

2711240741
00

0
14713

14713

00
1003317509

1003317509

00
66028519

66028519

10477
0

10477

8498
0

0
8498

8498

00000
42379359070206919905257557

1852015073
00

1852015073

169439342
00

169439342

0
9229

9229

3231335727
00

1782623509

1448712218

2221621632
00

2221621632

0
10546

10546

5025489709562
000

5025489709562

00000
125563354832069199057831

125563354832069199057831

00
1200331877

1200331877

1140312294
00

1140312294

0
5997

5997

720581461739042
000

720581461739042

51417
0

51417

00
3484217763

3484217763

7903
0

7903

21605
0

6522
0

6522

0
15083

15083

0000
1112543887215086105022

0
15332

15332

4983
0

4983

16450
0

16450

818912455
00

818912455

2020124796
00

2020124796

12413
0

12413

0000
82864388721508618593

82864388721508618593

0
7607

0
7607

7607

00
1423431848

00
1423431848

971712520
00

9717
0

9717

12520
0

12520

00
451719328

451719328
00

451719328

0000
109449334151171871905

1000021171871905
000

1000021171871905
000

33072
0

33072
0

33072

00
972915598

00
972915598

972915598

00
1835911450

11450
0

11450

18359
0

18359

719141171811785
000

0
12028

12028

10167
0

10167

6201
0

6201

11945
0

11945

315731171811785
000

315731171811785

00
944733415

9447
0

0
9447

9447
0

9447

33415
0

0
33415

33415
0

33415

29780954139378990596885548118813517
000000

26165143969746975934707391557297252
000000

0
12033

0
12033

12033
0

12033

00
1202131622

0
12021

12021
0

12021

0
31622

31622
0

31622

00
57386256297

0
57386

0
57386

57386

0
256297

0
232451

208243

24208

23846
0

23846

0
14424

0
14424

14424
0

14424

000000
2568518327567697593470796091297252

00
123249499

00
123249499

123249499

2556194327567697593470796091287753
000000

709903963125057
000

709903963125057

00
1843014346

11263

716714346

0
15776

15776

0
27707

27707

000000
4996996763822146197892059698129

4996996763822146197892059698129

32754
0

32754

14023
0

14023

15790229529
00

15790229529

0000
110584226441485031613

110584226441485031613

000000
2667526499415771149181442290475

2667526499415771149181442290475

0
13784

13784

24341
0

24341

0
17144

17144

0
20767

20767

3626012277
00

3626012277

124006310313110467
000

124006310313110467

000
412541699214426

412541699214426

00
215397547

00
215397547

61237547
00

61237547

0
7516

7516

0
7900

7900

361581169632014662178156561516265
000000

17659276691517032914
0000

17659276691517032914
0000

00
1765920208

1765920208

000
276691517012706

276691517012706

0000
3717897798193041664

180219779
00

00
82239779

82239779

0
9798

9798

0
81930

81930
0

81930

00
1915741664

00
1915741664

1915741664

22571616963201462473059461165388
000000

00
1386713440

00
1386713440

1386713440

21184916963201462473059461151948
000000

0000
4005720146936226496

4005720146936226496

10392
0

10392

0
8727

8727

0
8632

8632

0
12163

12163

13079
0

13079

173061536813587
000

173061536813587

0000
22009169631018412257

22009169631018412257

00
991410762

991410762

00
212248588

212248588

00
1114913768

1114913768

00
2275969790

2275969790

87868883
00

87868883

0
12453

12453

000
1437991946720

1437991946720

00
46880191285

0
19980

19980
0

19980

48720
0

18061
0

18061

0
11863

11863

0
18796

18796

0
6918

6918
0

6918

0
5785

0
5785

5785

00
1003412985

1003412985
00

1003412985

29928103815
00

42311
0

42311

0
7169

7169

0
24119

24119

0
8630

8630

731510204
00

731510204

00
1398320012

1398320012

3414885014
00

1098956240
00

20277
0

20277

00
1098914241

1098914241

0
11118

11118

10604
0

10604

0
6512

0
6512

6512

00
2315922262

972411960
00

972411960

1343510302
00

1343510302

0
14457

0
14457

0
14457

0
10963

10963
0

10963

0
3494

0
3494

3494

5379
0

0
5379

0
5379

0
5379

0
5379

5379

17214
0

0
17214

17214
0

0
17214

0
17214

17214

8702
0

8702
0

8702
0

8702
0

0
8702

8702

12603
0

0
12603

0
12603

12603
0

0
12603

12603

0
5897

5897
0

0
5897

0
5897

0
5897

5897

5969
0

0
5969

5969
0

5969
0

5969
0

5969

10196
0

10196
0

0
10196

0
10196

0
10196

10196

8827
0

0
8827

8827
0

8827
0

0
8827

0
8827

8827

2257138279
00

2257138279
00

00
2257138279

00
2257118247

00
2257118247

00
1099118247

1099118247

0
11580

11580

10047
0

0
10047

0
10047

10047

9985
0

0
9985

0
9985

9985

15734
0

15734
0

15734
0

0
15734

0
15734

15734
0

15734

375754028840
000

0
3757

3757
0

0
3757

0
3757

0
3757

3757

3483
0

0
3483

3483
0

3483
0

0
3483

3483

5357
0

0
5357

0
5357

0
5357

0
5357

5357

5402
0

0
5402

5402
0

5402
0

5402
0

5402

0000000
16443535107326398804289501711914472985891423601

0000000
1385949258976075495151272524202815149405589

000
195171530015150

000
195171530015150

00
1277615300

1277615300
00

1277615300

0
15150

0
15150

15150

0
6741

6741
0

6741

745111648
00

745111648
00

0
7451

7451
0

7451

0
11648

11648
0

11648

226121073311165310919702301514974935
0000000

0000000
226121073311165310919702301514974935

11648
0

0
11648

11648

2261295595
00

2261239948
00

2261239948

0
55647

55647

11736116531091911416151497969
000000

000000
11736116531091911416151497969

11736116531091911416151497969

0
8293

0
8293

8293

5881447025
00

23699
0

23699

23326
0

23326

0
58814

58814

000
5908951145023585

00
1996323585

1996323585
00

0
23585

23585

0
19963

19963

22554511450
00

00
22554511450

00
22554511450

22554511450

0
16572

0
16572

0
16572

16572

00000
3263471531147300623505445923

0
6695

6695
0

6695
0

6695

00000
3196521531147300623505445923

83423
0

83423
0

83423

0
5719

5719
0

5719

0
19504

19504
0

19504

3001481407951300623505440204
00000

0
33230

33230

0
28934

28934

0
30942

30942

35054
0

35054

40204
0

40204

0
72872

72872

30062
0

30062

0
107384

107384

0
54846

34336

20510

448121335079
00

448121335079

0
39773

39773
0

39773

0
9134

9134
0

0
9134

9134
0

9134

0
21062

0
21062

0
21062

21062
0

21062

134733776062709
000

00
3776036529

36529
0

0
36529

36529

0
37760

0
37760

37760

13473
0

0
13473

13473
0

13473

26180
0

0
26180

26180
0

26180

896377226630638421464293121444162505
000000

896377226630638421464293121444162505
000000

000
193612032450742

0
17280

17280

193612032433462
000

193612032433462

5689215454
00

0
15483

15483

0
15090

15090

0
13588

13588

1273115454
00

1273115454

82012422663063842144396912144475055
000000

0
16635

16635

000000
1194403734633142144396912144440188

1194403734633142144396912144440188

774431143633070018704
0000

774431143633070018704

56877674921
00

56877674921

0
16163

16163

0
37830

37830

0
21254

0
21254

21254

00
27472175442

27472175442
00

27472175442
00

0
175442

175442

0
27472

27472

000
509291418762727

000
509291418762727

509291418762727
000

509291418762727
000

000
509291418762727

509291418762727

000
1348683928953

1348683928953
000

000
1348683928953

1348683928953
000

1348683928953
000

1348683928953

0000000
15472175704229250124183288357212598467314

0
6586

0
6586

6586
0

6586
0

6586

00
782457527

00
782457527

00
782430613

30613
0

30613

7824
0

7824

26914
0

26914
0

26914

27689
0

0
27689

0
27689

0
10329

10329

0
17360

17360

000000
321183687928357794137886311985

000000
321183687928357794137886311985

96721945911985
000

0
11985

11985

00
967219459

967219459

3211836879283573497359404
00000

0
11704

11704

36879283572710441901
0000

36879283572710441901

000
20414786917503

20414786917503

34768
0

34768
0

34768

0000000
1547214358619237189240173431133735397802

0
85095

0
43808

43808
0

43808

41287
0

41287
0

41287

3518084005409326113274394101599
000000

3518084005409326113274394101599
000000

351802921117433212333598679387
000000

351802921117433212333598679387

31803234992259538408
0000

31803234992259538408

000
229911730422212

229911730422212

0
14134

14134
0

0
14134

14134

723951083664830811229959341167202
000000

0000
64069331894574459341

0000
64069331894574459341

64069331894574459341

72395442971511966555167202
00000

24768
0

24768

000
7239533159142434

7239533159142434

000
232661511915699

232661511915699

2103117697
00

2103117697

5967
0

5967
0

0
5967

5967

000
95053601129772

000
95053601129772

95053601129772
000

95053601129772

31808
0

31808
0

31808
0

31808
0

0
31808

31808

0
8880

8880
0

0
8880

0
8880

8880
0

8880

0000000
183123232767580573125810965267061889456163

5178342257299985558420865
00000

13697
0

0
13697

0
13697

13697

00000
517834225729998555847168

0
12776

0
12776

12776

0000
51783422572999842808

000
131291298213842

131291298213842

13030
0

13030

1365621762
00

1365621762

0000
14247204951701615936

14247204951701615936

0
10751

10751

7168
0

0
7168

7168

000000
60567810200494473191030218842

00000
1020049447319103027305

00000
1020049447319103027305

00000
1020049447319103027305

1020049447319103027305

00
60567811537

0
605678

605678
0

605678

11537
0

11537
0

11537

2783442278
00

0
42278

0
42278

0
42278

42278

0
27834

0
27834

27834
0

27834

103506163746270373122316758976751587416456
0000000

0000000
86547901395832430861405974037918715

0
8654

8654
0

8654

12839
0

0
12839

12839

0
790139

790139
0

790139

00000
5832430861277584037918715

0
13308

13308

4501630861277584037918715
00000

4501630861277584037918715

000000
175402659251047079671611208275004

00
134858168850

62955
0

62955

71903139531
00

71903139531

0
29319

29319

0
46679

0
46679

46679

47065
0

0
47065

47065

11208
0

11208
0

11208

0000
84388626775191159089

84388626775191159089
0000

843886267751911

59089

17540
0

17540
0

17540

4203044805
00

4203044805
00

4203044805

00000
88203324612049928222015

3324611176
00

00
3324611176

3324611176

882012049928210839
0000

882012049928210839
0000

882012049928210839

000
1715211858740288

0
118587

38340
0

38340

0
80247

80247

00
1715240288

1715240288
00

1715240288

00000
68492548152106116533386760434

000
21301212683848548

0
47613

47613

0
49013

49013

21301116057
00

21301116057

848548
0

848548

0
88174

88174
0

88174

0000
223091099752158742533

0000
22309872332158742533

22309872332158742533

22742
0

22742

0
24882

0
24882

24882

0
17901

17901
0

17901

000
137320212617312280

000
89931212617312280

89931212617312280

0
47389

47389
